# Binding of guide piRNA triggers methylation of the unstructured N-terminal region of Aub leading to assembly of the piRNA amplification complex

**DOI:** 10.1101/2020.07.14.203323

**Authors:** Xiawei Huang, Hongmiao Hu, Alexandre Webster, Fan Zou, Jiamu Du, Katalin Fejes Toth, Alexei A. Aravin, Sisi Li

## Abstract

Piwi proteins use guide piRNAs to repress selfish genomic elements, protecting the genomic integrity of gametes and ensuring the fertility of animal species. Efficient transposon repression depends on amplification of piRNA guides in the ping-pong cycle, which in Drosophila entails tight cooperation between two Piwi proteins, Aub and Ago3. Here we show that post-translational modification, symmetric dimethylarginine (sDMA), of Aub is essential for piRNA biogenesis, transposon silencing and fertility. Methylation is triggered by loading of a piRNA guide into Aub, which exposes its unstructured N-terminal region to the PRMT5 methylosome complex. Thus, sDMA modification is a signal that Aub is loaded with piRNA guide. Amplification of piRNA in the ping-pong cycle requires assembly of a tertiary complex scaffolded by Krimper, which simultaneously binds the N-terminal regions of Aub and Ago3. To promote generation of new piRNA, Krimp uses its two Tudor domains to bind Aub and Ago3 in opposite modification and piRNA-loading states. Our results reveal that post-translational modifications in unstructured regions of Piwi proteins and their binding by Tudor domains that are capable of discriminating between modification states is essential for piRNA biogenesis and silencing.

**Highlights:**

- sDMA modification of Aub is essential for ping-pong cycle, transposon silencing and fertility.
- piRNA loading triggers Aub sDMA modification by exposing its unstructured N terminal region to the methylosome complex.
- A single molecule of Krimp simultaneously binds piRNA-bound Aub and empty Ago3 to assemble ping-pong complex.
- The distinct structure of two Tudor domains of Krimp ensures binding of Ago3 and Aub in opposite modification and piRNA-loading states.

## Introduction

The PIWI-interacting RNA (piRNA) pathway acts as a conserved defensive system that represses the proliferation of transposable elements (TEs) in the germline of sexually reproducing animals (Kalmykova, Klenov, and Gvozdev 2005; Brennecke et al. 2007; Li et al. 2009; Aravin et al. 2001; Gunawardane et al. 2007; Nishida et al. 2007; Vagin et al. 2006). Loss of Piwi proteins causes derepression of transposons associated with gametogenesis failure and sterility in flies and mice (Khurana and Theurkauf 2010; Cox et al. 1998; Girard et al. 2006; Carmell et al. 2007). Piwi proteins recognize transposon targets with help of the associated small (23-30 nt) non-coding RNA guides, piwi-interacting RNAs.

Piwi proteins belong to the conserved Argonaute protein family present in all domains of life (Cox et al. 1998; Carmell et al. 2002; Cox, Chao, and Lin 2000). Argonautes bind nucleic acids guides and share common domain architecture, all containing the conserved N, PAZ, MID and PIWI domains (Wei et al. 2012; Song et al. 2004; Elkayam et al. 2012; Schirle and MacRae 2012). The MID and PAZ domains bind the 5’ and 3’ ends of the guide RNA, respectively (Boland et al. 2011; Frank, Sonenberg, and Nagar 2010; Ma, Ye, and Patel 2004; Song et al. 2003; Song et al. 2004; Yan et al. 2003). The PIWI domain contains an RNase-H like fold with a conserved amino acid tetrad that endows Argonautes with endonuclease activity for precise cleavage of the target (Parker, Roe, and Barford 2004; Song et al. 2004; Martinez and Tuschl 2004). The degradation of complementary target mRNA by Piwi proteins can trigger the generation of new RNA guides in a process termed the ping-pong cycle. Ping-pong requires cooperativity between two Piwi molecules as the product resulting from target cleavage by one protein is passed to the other and is converted to a new piRNA guide. In *Drosophila*, two distinct cytoplasmic Piwi proteins, Aub and Ago3, cooperate in the ping-pong cycle with each protein generating an RNA guide that is loaded into its partner. Amplification of piRNA guides through the ping-pong cycle is believed to be essential for efficient transposon repression as it allows the pathway to mount an adaptive response to actively transcribed transposons (Brennecke et al. 2007; Gunawardane et al. 2007; Huang, Fejes Toth, and Aravin 2017).

In addition to four conserved domains, eukaryotic members of the Argonaute family, including Piwi proteins, contain a N-terminal extension region of various lengths with low sequence conservation. Structural studies of Agos suggest that the N-terminal regions adopt a disordered conformation (Faehnle et al. 2013; Nakanishi et al. 2012; Schirle, Sheu-Gruttadauria, and MacRae 2014; Park et al. 2019; Schirle and MacRae 2012). Despite low overall conservation, the N-terminal region of the majority of Piwi proteins harbors arginine rich (A/G)R motifs. In both insects and mammals these motifs were shown to be substrates for post-translational modification by the PRMT5 methyltransferase, which produces symmetrically methylated arginine (sDMA) residues (Vagin, Hannon, and Aravin 2009; Honda, Kirino, and Kirino 2014; Kirino et al. 2009; Reuter et al. 2009; Nishida et al. 2009). Loss of *Prmt5* (encoded by the Csul and Vls genes) in *Drosophila* leads to reduced piRNA level and accumulation of transposon transcripts in germ cells, suggesting that sDMA modification of Piwis plays an important role in the piRNA pathway.

Multiple Tudor domain-containing proteins can bind to sDMA modifications. Aromatic residues in binding pocket of Tudor domains form cation-π interactions with sDMA (Tripsianes et al. 2011; Liu, Wang, et al. 2010; Liu, Chen, et al. 2010; Chen et al. 2011). Studies in *Drosophila* and mouse revealed that several Tudor domain-containing proteins (TDRDs) interact with Piwis and are involved in piRNA-guided transposon repression, although their specific molecular functions remains poorly understood (Nishida et al. 2009; Siomi, Mannen, and Siomi 2010; Chen et al. 2011; Handler et al. 2011; Sato, Iwasaki, Siomi, et al. 2015). Previously we found that the Tudor-domain containing protein Krimper (Krimp) is required for ping-pong piRNA amplification and is capable of both self-interactions and binding of the two ping-pong partners, Aub and Ago3 (Webster et al. 2015). Krimper co-localizes with Aub and Ago3 in nuage, a membraneless perinuclear cytoplasmic compartment where piRNA-guided target degradation and ping-pong are proposed to take place. Ago3 requires Krimper for recruitment into this compartment, though Aub does not. These results led to proposal that Krimp assembles a complex that brings Ago3 to Aub and coordinates ping-pong in nuage (Webster et al. 2015). However, the architecture of the ping-pong piRNA processing (4P) complex and the extent to which Krimp regulates ping-pong remained unresolved.

Both the ping-pong cycle and sDMA modification of Piwi proteins are conserved features of the piRNA pathway, found in many organisms, suggesting that these processes are essential for pathway functions (Honda, Kirino, and Kirino 2014; Siomi, Mannen, and Siomi 2010; Brennecke et al. 2007; Aravin et al. 2008; Gunawardane et al. 2007). sDMA modification of Piwis provides a binding platform for interactions with Tudor-domain proteins, however, its biological function and regulation are not known. Despite the essential role of ping-pong in transposon repression, we similarly have little understanding of the molecular mechanisms that control this process. Here we revealed the biological function of Aub and Ago3 sDMA modifications and show that it plays an essential role in orchestrating assembly of the 4P complex in the ping-pong cycle. The modification signals whether Piwi proteins are loaded with guide piRNA, and this information is used to assemble a ping-pong complex that is receptive for directional transfer of RNA to an unloaded Piwi protein.

## Results

### Krimper can simultaneously bind Aub and Ago3 with specificity for methylation state using two separate Tudor domains

The essential step in the ping-pong cycle is the transfer of RNA between two Piwi proteins, Aub and Ago3. This process requires that the two proteins are in physical proximity to each other. Previously we found that Krimp binds both Aub and Ago3 suggesting that it might bring them into physical proximity to allow transfer of the RNA during ping-pong (Webster et al. 2015). This might be possible through self-interaction of two Krimp molecules, each binding one Piwi protein, or by a single Krimper molecule simultaneously interacting with both Aub and Ago3. To discriminate between these two models, we employed two-step sequential immunoprecipitation to examine complex formation between Aub, Ago3 and a Krimper that lacks the N-terminal self-interaction region (residues 1-301aa) (Fig.1A and C). We immuno-precipitated tagged Aub followed by elution of purified complexes and another round of immunoprecipitation against the self-interaction-deficient Krimper. If a single Krimper molecule is capable of simultaneously interacting with Aub and Ago3, then the final immunoprecipitated complexes should contain Ago3 in addition to Krimper and Aub. All three proteins were detected by Western blot after sequential purification (Fig.1B). This indicates that a single Krimper molecule can concomitantly interact with both Aub and Ago3.

**Figure 1.**
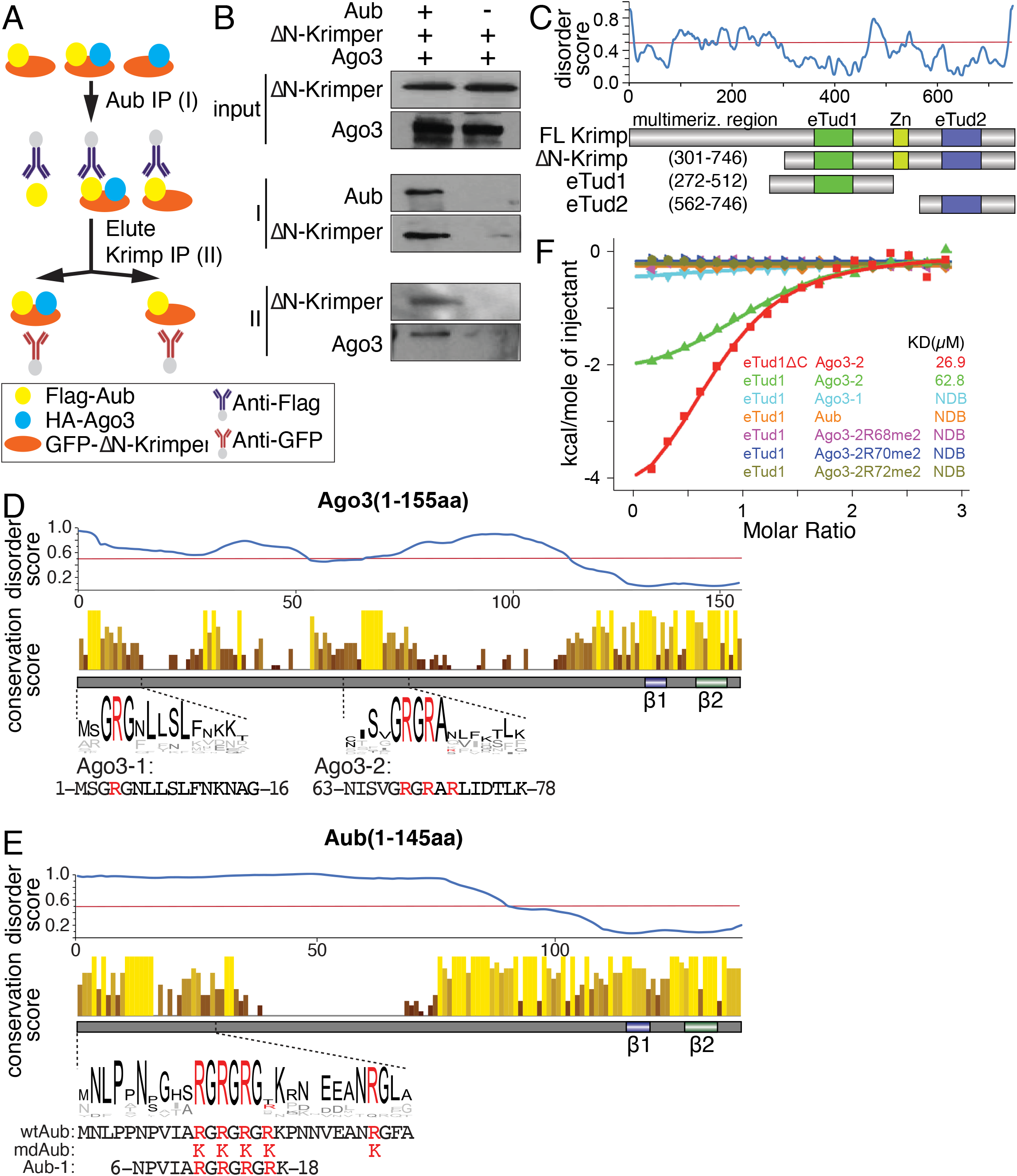
Krimper monomer simultaneously binds N-terminal regions of Aub and Ago3. (A) Scheme depicting two-step sequential immunoprecipitation. FLAG tagged Aub, HA tagged Ago3 and GFP tagged truncated Krimper lacking its N terminal dimerization domain were co-expressed in S2 cell. FLAG-immunoprecipitation was followed by elution and GFP IP. (B) Krimper monomer interacts with Aub and Ago3 simultaneously. Western blot detection of FLAG-Aub, HA-Ago3 and GFP-deltaN-Krimp in input and after 1^st^ FLAG- and 2^nd^ GFP-IP. (C) Top: Protein disorder prediction of Krimper. Bottom panel: Domain architecture of Krimp and constructs used for ITC and structural analyses. (D) Conservation analysis of Ago3 (1-155aa). Top: Disorder prediction of Ago3 (1-155aa). Middle: Conservation score of each amino acid within Ago3(1-155aa). Conserved RG-repeat regions are. Bottom: Peptides used for ITC and structural analysis. (E) Conservation analysis of Aub (1-145aa). Top: Disorder prediction of Aub (1-145aa). Middle: Conservation score of each amino acid within Aub(1-145aa). Conserved RG-repeat region is enlarged. Bottom: Sequence difference between wtAub and mdAub. Five arginines within N terminus were replaced by lysines. Aub-1 peptide used for ITC and structural analysis. (F) eTud1 preferentially binds unmethylated Ago3. ITC analysis of Krimper eTud1 with different Ago3 or Aub peptides.

To understand how Krimper is able to bind Aub and Ago3 simultaneously, we analyzed its domain architecture. In addition to its self-interacting N-terminal region, Krimper contains two extended Tudor domains: eTud1(272-512) and eTud2 (562-746) (Fig. 1C). Tudor domains were previously implicated in protein-protein interactions through specific binding of methylated Arginine or Lysine residues (Liu, Wang, et al. 2010; Liu, Chen, et al. 2010; Tripsianes et al. 2011). Piwi proteins, including Aub and Ago3, harbor conserved symmetrically dimethylated Arginine (sDMA) residues embedded into RA/RG-motifs in their N-terminal extended regions (Kirino et al. 2009; Kirino et al. 2010; Nishida et al. 2009) In agreement with published structures of other Piwi and Ago proteins, which suggest that N-terminal extensions of these proteins adopt an unstructured conformation (Faehnle et al. 2013; Matsumoto et al. 2016; Nakanishi et al. 2012; Schirle and MacRae 2012; Yamaguchi et al. 2020; Elkayam et al. 2012), the N-terminal extended regions of both Aub and Ago3 have high levels of predicted disorder (Fig. 1D and 1E). Remarkably, RA/RG motifs, which are subject to sDMA modification are the only conserved sequences in the N-terminal regions of Aub and Ago3. The N-terminal regions are otherwise highly variable even between closely related Drosophila species (Fig. 1D and 1E). We previously found that the Krimper/Aub interaction depends on Aub methylation and found that the eTud2 domain of Krimper specifically binds sDMA-modified Aub (Webster et al. 2015). However, in contrast to Aub, Ago3 was shown to interact with Krimper in its unmethylated state (Webster et al. 2015; Sato, Iwasaki, Shibuya, et al. 2015), raising the question of how these two proteins interact.

Previous studies revealed that the N-terminal (1-83aa) sequence of Ago3 is both necessary and sufficient for Krimper binding (Sato, Iwasaki, Shibuya, et al. 2015). However, the region of Krimper responsible for Ago3 binding remained unknown (Webster et al. 2015). To study the Ago3-Krimp interaction, we expressed and purified eTud1 and eTud2 domains of Krimper (Fig. 1C) and tested their binding to peptides derived from the N-terminal Ago3 sequence using isothermal titration calorimetry (ITC) (Fig. 1D). We found that eTud1 binds to the Ago3-2 peptide (aa residues 63-78 of Ago3, which includes the conserved RA/AG motif) with a binding affinity of 62.8 μM (Fig. 1F). Importantly, eTud1 does not detectably interact with the Aub peptide that binds eTud2 (Fig 1F). Thus, the two Tudor domains of Krimper have different binding preferences ensuring that Aub and Ago3 can bind Krimper simultaneously without direct competition for a binding site.

The Aub-Krimper (eTud2) interaction requires methylation of at least one of five Arg residues positioned within the RA/RG motif of the Aub N-terminal region (Webster et al. 2015). Remarkably, the Ago3-2 peptide that binds eTud1 domain also contains three Arg residues (R68, 70 and 72) within the RG/RA motif (Fig. 1D), which can be methylated *in vivo*, however, methylation is not required for Ago3 binding to Krimp (eTud1) (Fig. 1F). In fact, in agreement with *in vivo* experiments that indicate that Ago3 is bound to Krimp in its unmethylated state (Sato, Iwasaki, Shibuya, et al. 2015; Webster et al. 2015), methylation of any of the three Arg residues disrupts binding of the Ago3-2 peptide to eTud1 (Fig. 1F). Overall, our results indicate that binding of both Aub and Ago3 to Krimp is mediated by the RG/RA-motifs embedded in their unstructured N-terminal extended regions. However, the two proteins interact with Krimp in opposite modification states: Aub must be sDMA-modified, while Ago3 remains unmethylated. The distinct specificities of the two Tudor domains explain how Krimper can simultaneously bind Aub and Ago3 to bring them into physical proximity for ping-pong.

### Structural differences in the two Tudor domains of Krimper explain their different binding preference towards Arg-modified Aub and unmodified Ago3

To gain a detailed understanding of the interaction between Krimp (eTud2) and Aub, we undertook structural studies. We solved the crystal structure of the Krimp (eTud2) domain in complex with the Aub-R15me2 peptide that showed highest binding affinity in ITC experiments (Webster et al. 2015). In the 2.7 Å structure the eTud2 domain adopted a typical Tudor subdomain and a staphylococcal nuclease (SN)-like subdomain (Tudor-SN) architecture (Fig. 2B and 2C). sDMA-modified Arg-containing peptide binds within a negatively-charged cleft between the Tudor and SN domains, and is specifically recognized by a conserved aromatic cage consisting of four aromatic amino acids Tyr624, Tyr630, Tyr646 and Tyr649 (Fig. 2A, 2D and 2E), stabilized further by hydrophobic and cation-π interactions (Fig. 2D). In addition, eTud2 Asn651, which is conserved amongst other extended Tudor domains, forms a hydrogen bond with the guanidyl nitrogen of the methylated Arg15 (Fig. 2D and S2A). Besides Arg15, other flanking arginine residues in the (G/A)R motif of Aub also contribute to protein-peptide binding, (Fig. S2A). Together, these structural features of eTud2 explain its specific binding to sDMA-modified Aub.

**Figure 2.**
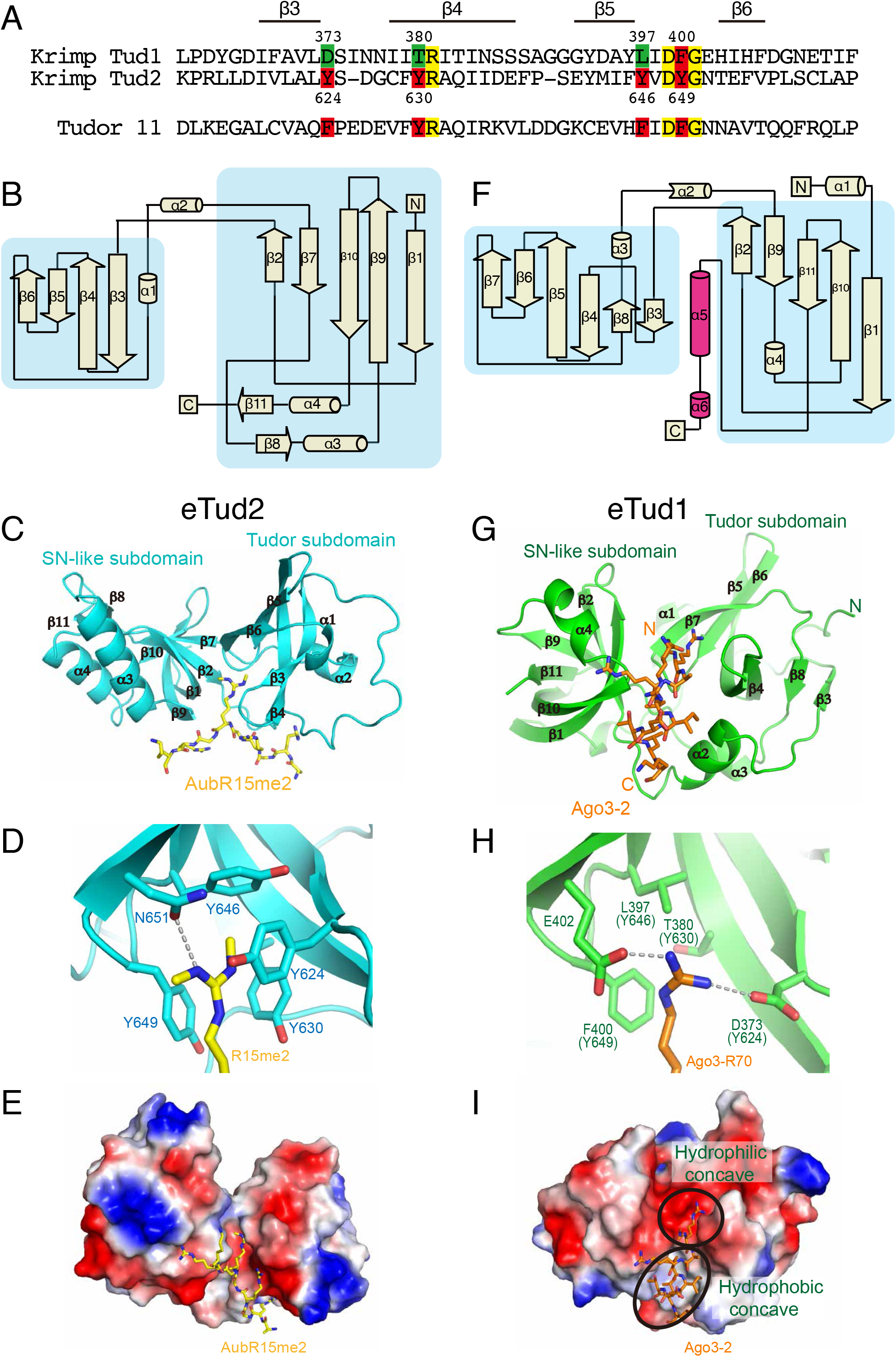
Structure of Krimper eTud2-Aub and eTud1-Ago3 complex. (A) Krimper contains the canonical eTud2 domain and the non-canonical eTud1 domain. Alignments of Krimper Tudor domains with Tudor1 domain 11 are shown. Residues forming the me-Arg binding pocket (red) and corresponding residues in tud1 (green) are shown. Additional conserved residues marked yellow. Structural features are shown above. (B) Overall topology demonstration of Krimper eTud2 domain. The Tudor and SN-like subdomain are indicated. (C) Overall structure of the Krimper eTud2-Aub-R15me2 complex with the bound Aub-R15me2 peptide shown in yellow. (D) Enlarged view of the eTud2 aromatic cage. (E) Electrostatic surface of eTud2 with bound Aub-R15me2 peptide shown in yellow. (F) Overall topology demonstration of Krimper eTud1 domain. The Tudor and SN-like subdomain are indicated. Latch helix is marked magenta. (G) Overall structure of the Krimper eTud1-Ago3-2 complex with bound Ago3-2 peptide shown in orange. (H) Enlarged view of eTud1 binding pocket with inserted Ago3 R70 residue indicated in orange. (I) Electrostatic surface of eTud1 with bound Ago3-2 peptide shown in orange.

Unlike eTud2, Krimp (eTud1) has an unusual binding specificity towards the unmethylated Ago3 sequence containing the (G/A)R motif. To elucidate the interaction between eTud1 and Ago3 we solved the crystal structure of eTud1 in both apo and peptide bound forms. In the 2.1 Å eTud1 apo structure, crystals belong to the *P*2_1_2_1_2_1_ space group with two monomers in the asymmetric unit (Table S1). The overall structure is similar to eTud2 apart from the C-terminal region (484-511aa) that forms a long α-helical fold that inserts into the narrow interface between the Tudor and SN-like subdomains. We named this novel C-terminal helical topology the ‘latch helix’ (Fig. 2F and S2B). The latch helix is stabilized through formation of extensive hydrogen bonding and salt bridge interactions with its amphipathic concave binding site within eTud1. Detailed interaction patterns and paradigm are shown in Fig. S2F.

Next, we determined the crystal structure of eTud1-Ago3-2 peptide complex at 2.4 Å resolution (Table S1). The structure of eTud1 in the eTud1-Ago3-2 peptide complex exhibits almost an identical fold to the apo form, with an overall root mean square deviation of 0.60 Å (Fig. S2D). However, the Ago3 peptide displaces the ‘latch helix’ and adopts an α-helical structure to bind in the same cleft between the Tudor and SN-like subdomains, but with reverse directionality compared to the ‘latch helix’ (Fig. 2G and S2C). These results suggest that the ‘latch helix’ might regulate binding of Ago3 to eTud1. Indeed, ITC results showed that eTud1 devoid of the latch helix (named eTud1 ΔC) has a higher binding affinity towards the unmethylated Ago3-2 peptide (Fig 1F).

The Ago3 peptide binds both a hydrophilic concave and a hydrophobic concave region in the cleft between the Tudor and SN-like subdomains of eTud1 (Fig. 2I). However, instead of a conserved aromatic cage that interacts with sDMA in other canonical Tudor domains including eTud2, the binding of unmethylated Arg70 of Ago3 in eTud1 is provided by a mostly hydrophilic pocket composed of three non-aromatic amino acids (Asp373, Thr380 and Leu397) (Fig. 2H and 2I). Arg70 forms hydrogen bonds with Asp373 and Glu402 in eTud1, supplemented by a cation-π interaction with Phe400 (Fig. 2H). Many extended Tudor domains adopts an incomplete aromatic cage, which may prefer unmethylated arginine peptides. In human TDRD2 protein, replacement of one of the four aromatic amino acids in the aromatic cage by a leucine residue, results in the preferential binding of unmethylated (A/G)R repeat with higher affinity (Zhang et al. 2017). In addition, the backbone of Arg68 and the side chain of Arg72 form hydrogen bonds with Asn376 and Thr445, respectively (Fig. S2E). Notably, the side chain of Phe400 is rotated by 90° to form a stronger cation-π interaction with Arg70 when compared with the structure in the apo state (Fig. S2G). Adjacent to the hydrophilic concave region there exists a hydrophobic concave region formed by Phe400, Trp339, and Tyr446. The Ago3 peptide forms a hydrophobic contact with the hydrophobic concave region utilizing Leu73 and Leu77 that strengthens its interactions (Fig. 2H and S2E). The bound unmethylated arginine peptide in the Krimper eTud1 complex adopts the same directionality and similar localization as its counterpart in the TDRD2 complex (PDB code: 6B57) (Fig. S3A and S3B). However, the peptide-binding interface of eTud1 is much narrower compared with the extended and negatively charged interface of TDRD2. Thus, eTud1 specifically recognizes unmodified R(A/G) motif in Ago3 using a unique hydrophilic concave and a hydrophobic concave region in the interface between the Tudor and SN-like subdomains.

Although the overall topology of eTud1 is similar to other extended Tudor domains, eTud1 has its own unique folding and recognition characteristics. First, the N-terminus of eTud1 contains a long-loop (spanning amino acids 272-300) connecting and interacting with both the Tudor and the SN-like subdomains (Fig. S3C). Pro292 within this N-terminal loop forms a CH-π interaction with Tyr365, Glu297 in the loop region forms a hydrogen bonding with the backbone of Ser387, Arg300 in the loop forms a hydrogen bond and salt bridge interactions with Asp394, while Tyr304 in the loop forms cation-π interaction with Arg428 in the SN-like domain. All these intramolecular interactions help to stabilize the overall structure of eTud1 (Fig. S3C). Second, the linker helix α2 is shorter and shifted towards the Tudor-SN interface compared with other extended Tudor domains (PDB code: 3OMC), facilitating the interactions with the C-terminal ‘latch helix’ or bound Ago3-2 peptide (Fig.S3D).

Overall, our results reveal that the two Tudor domains of Krimper have distinct architectures that are responsible for differential binding of two Piwi proteins. Aub and Ago3 have similar organization, with N-terminal (G/A)R motifs that are subject to sDMA modification *in vivo*. However, while Aub interacts with the eTud2 domain in a methylated state through sDMA binding to a conserved aromatic cage, Ago3 binds the eTud1 domain in an unmethylated state employing a binding pocket that is distinct from other Tudor domain interactions. Furthermore, eTud1 possesses a ‘latch helix’ that must move out of the cleft to allow Ago3 binding providing a potential regulatory mechanism. These results highlight the crucial role that sDMA modifications of piwi proteins play in regulating the formation of the tertiary ping-pong complex.

### sDMA modification of Aub is required for piRNA biogenesis through the ping-pong cycle but is dispensable for loading of piRNA into Aub

Alongside previous studies, our results demonstrate the critical function that the methylation states of Aub and Ago3 plays in their binding to Krimper. In order to investigate the function of sDMA modification of Aub in the broader piRNA pathway *in vivo*, we generated transgenic flies expressing a <u>m</u>ethylation-<u>d</u>eficient version of Aub (mdAub) by mutating five Arg into Lys in the N-terminal region (Fig. 1E). The methylation of wild-type Aub was readily detected using Western blotting with SYM11 antibody that recognizes sDMA residues (Fig. 3A). In contrast, mdAub produced no signal, indicating that it indeed lack sDMA modifications. Transgenic mdAub and control wild-type protein were expressed in the ovary on a similar level. Wild-type Aub localized to nuage, the membrane-less subcellular compartment that surrounds the nucleus of nurse cells and is believed to play an essential role in piRNA biogenesis and TE repression. mdAub is also recruited to nuage, although compared to the wild-type protein a higher fraction is dispersed in the cytoplasm (Fig. 3B, 3C and S4A). FRAP experiments show a slightly increased mobility of mdAub in nuage compared to wild-type Aub (Fig. 3C and S4B). Thus, the methylation status of Aub does not affect protein stability and has only a minor effect on subcellular protein localization.

**Figure 3.**
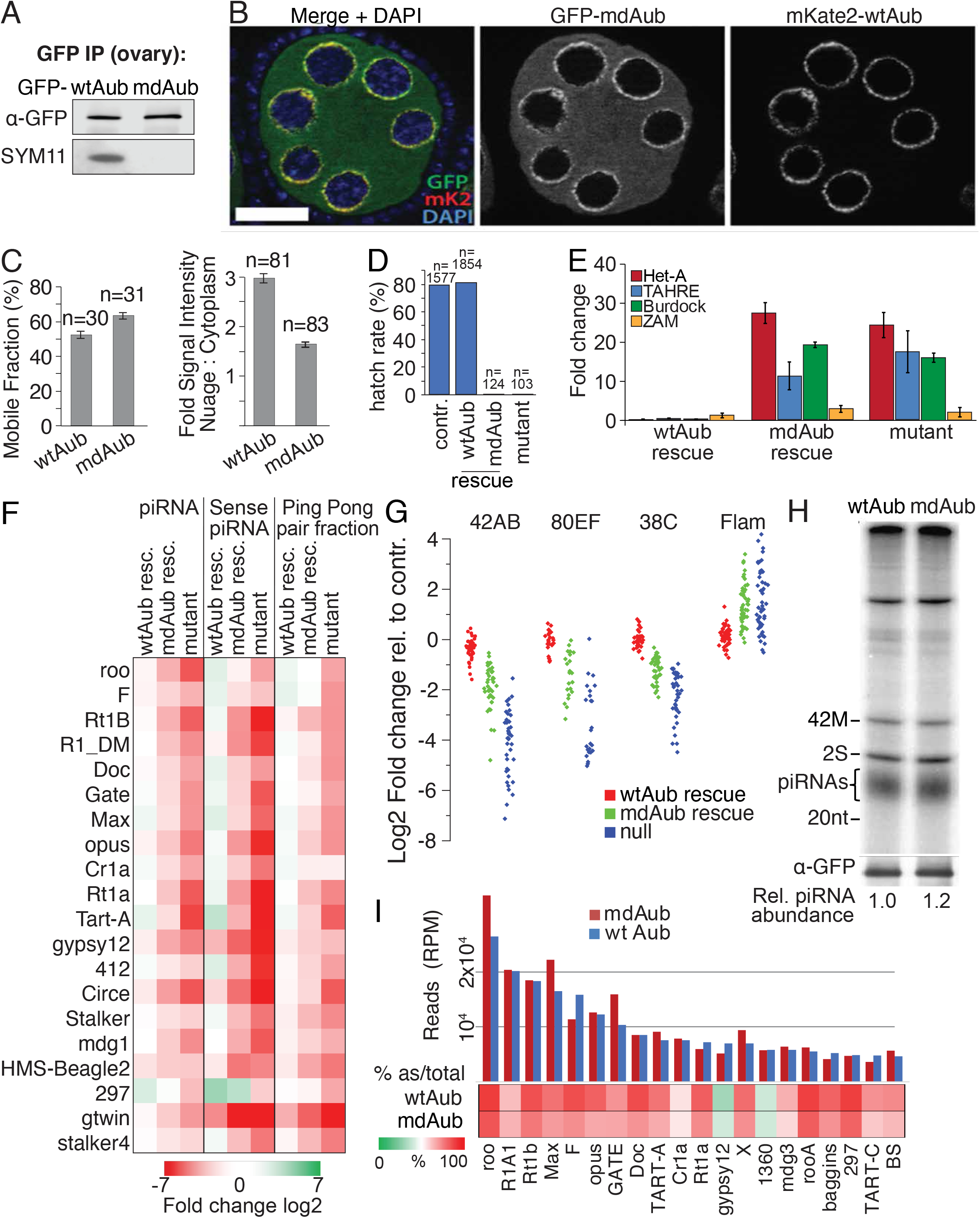
Arginine methylation of Aub is dispensable for localization and piRNA binding but required for ping-pong and TE repression. (A) Mutating conserved N-terminal Arginines to Lysines (mdAub) leads to loss of symmetric dimethylation of Aub. Similar amount of wtAub and mdAub was immunoprecipitated from ovaries. Methylation was detected by Western blot using the SYM11 antibody. (B) In the wildtype background mdAub colocalizes with wildtype Aub in nuage. GFP-mdAub and mKate2-wtAub were co-expressed in ovaries under the control of the endogenous Aub promoter. Scale bar: 5μm. (C) Arginine methylation deficiency slightly increases the fraction of mobile and dispersed cytoplasmic Aub protein. Left: the mobile fraction of nuage-localized GFP-wtAub and GFP-mdAub was determined in replicate FRAP experiments (constructs as in Fig. S4). Right: microscopic quantification of the ratio of GFP signal originating from nuage versus cytoplasm is shown. Error bars indicate st. dev. All transgenes are expressed under the control of the endogenous Aub promoter. (D) Arginine methylation of Aub is required for fertility. Hatching rate of heterozygous control, Aub^HN/QC^ mutant females and mutant females rescued with wt- or mdAub transgenes is indicated as % of eggs laid. (E) Arginine methylation of Aub is required for TE suppression. Fold changes of different TE transcripts in ovaries of flies with the indicated genotypes compared to the heterozygous control as measured by RT-qPCR. n=3, error bars indicate st. dev. (F) Arginine methylation of Aub is required for ping-pong processing. Total piRNA abundance, Aub cleaved (sense) piRNA abundance and the fraction of ping-pong pairs with 10nt overlap was analyzed in heterozygous control, wtAub rescue, mdAub rescue and aub mutant flies. The heatmap shows the fold change compared to the heterozygous control for the top 20 most abundant TE families present in the control ovary. (G) Aub Arginine methylation is required for generation of piRNAs from dual strand but not for single strand piRNA cluster. Reads uniquely mapping to different piRNA clusters was determined in heterozygous control, wtAub rescue, mdAub rescue and aub mutant flies and plotted as the fold change compared to the het control for four clusters (42AB, 38C, 80EF, Flam). Dot represents 1kb windows with uniquely mapping piRNAs within the clusters. (H) Aub Arginine methylation is not required for piRNA binding. GFP-tagged mdAub and wtAub were immunoprecipitated from ovaries where the transgene was expressed in the wildtype background. Protein level were detected by WB, associated RNAs were isolated, radiolabeled and run on a Urea PAGE gel. 42 nt ssRNA (42M) was spiked into each IP sample to control for labeling efficiency and RNA loss during isolation. Relative piRNA abundance normalized to IP-ed protein is estimated based on band intensity. (F) Arginine methylation of Aub does not greatly affect the TE and antisense fractions of Aub-bound piRNAs. Bar chart shows normalized read counts (RPM) of wtAub and mdAub bound small RNAs mapped to the 20 TE families with post abundant piRNAs. Aub transgenes were expressed on the wildtype background. Heatmap shows the % of antisense reads mapping to each TE family.

To explore the function of sDMA modification of Aub *in vivo*, we studied the ability of mdAub to rescue defects observed in Aub mutant flies. We expressed transgenic wild-type and methylation-deficient versions of Aub under the control of the endogenous Aub promoter and used genetic crosses to introduce these transgenes into the aub^HN/QC^ mutant background. Aub^HN/QC^ females are completely sterile due to severe defects in oocyte axis specification and DNA damage, presumably caused by derepression of multiple transposon families (Cook et al. 2004; Klattenhoff et al. 2007). While the wild-type Aub protein was able to rescue transposon derepression and sterility of the *aub* mutant background, methylation-deficient Aub failed to rescue both phenotypes (Fig. 3D and 3E). The inability of mdAub to rescue sterility and transposon activation suggests that sDMA modification is essential for Aub function *in vivo*.

Sequencing of total small RNA libraries from ovaries of aub mutant females that express wild-type or methylation-deficient variants of Aub revealed that piRNAs mapping to different transposon families and major piRNA clusters were severely reduced in the aub mutant (Fig. 3F and 3G). While expression of methylation-deficient Aub increased piRNA levels compared to the aub^HN/QC^ mutant, this increase was smaller than that observed in flies rescued with wild-type protein.

We previously proposed that binding of Aub and Ago3 to Krimp assembles a ping-pong piRNA processing (4P) complex that is essential for the ping-pong cycle (Webster et al. 2015). To explore the role of Aub methylation in ping-pong we analyzed piRNA profiles of aub mutant flies rescued with either wild-type or mdAub to find signatures of ping-pong processing: pairs of piRNAs whose 5’ ends are complementary across 10 nt, and piRNAs that are sense relative to TE and have A residues in position 10. In the piRNA libraries isolated from ovaries of mdAub-rescued flies, piRNAs with ping-pong processing signatures were significantly reduced for almost all major families of transposons compared to those from the wild type Aub rescue (Fig. 3F), indicating defects in the ping-pong cycle.

To explore if disruption of piRNA biogenesis might be caused by an inability of mdAub to be loaded with piRNAs, we expressed the mdAub transgene in wild-type flies that also express endogenous wild-type Aub. These flies have normal fertility and transposon expression (data not shown) indicating that expression of mdAub does not induce a dominant-negative effect. Next, we purified mdAub and control wild-type Aub protein, and radiolabeled isolated RNAs associated with both proteins. Quantification of the signal showed that small RNA loading of wild type and mdAub is similar (Fig. 3H). Small RNAs associated with the two proteins have a similar length distribution (Fig. S4C). To explore if small RNA loaded in mdAub might be different from sequences loaded into wild-type Aub protein we cloned and analyzed small RNAs residing in both complexes. Similar to wild-type Aub protein, the majority of sequences in mdAub are antisense to various transposons families (Fig. 3I). Overall, we did not find any abnormalities in the amount and composition of piRNA loaded into mdAub when it was expressed in the wild-type Aub background. Thus, the methylation status of Aub plays no direct role in piRNA loading.

Taken together, our results indicate that sDMA modification of Aub is essential for piRNA biogenesis through piRNA amplification in the ping-pong cycle and hence the repression of transposons. However, sDMA modification is not required for loading of piRNA into mdAub if cells also express wild-type Aub that ensures functional piRNA biogenesis. This finding corroborates our earlier proposal that Aub methylation is required for the formation of the Krimper, Aub and Ago3 complex, which in turn is required for Ago3 loading (Webster et al. 2015). Overall, our results indicate that the sDMA modification of Aub has a crucial role in piRNA biogenesis and specifically in the ping-pong amplification cycle.

### piRNA loading induces Aub methylation

Our results demonstrate that sDMA modification of Aub is essential for an ability of the piRNA pathway to generate a repertoire of guide piRNAs that effectively target and repress transposons. To understand the molecular mechanism of sDMA modifications we decided to explore if loading of piRNAs into Aub affects its modification. We used flies that express a mutant version of Aub that is deficient in piRNA binding (pdAub) due to mutation in two conserved residues within the 5’ end binding pocket located in the MID domain (Webster et al. 2015). We expressed pdAub on a wild-type background to preserve an intact piRNA biogenesis pathway and tested the methylation status of Aub using SYM11 antibody that recognizes methylated Arginine residues. Remarkably, we found that methylation of pdAub is strongly reduced compared to wild-type protein (Fig. 4A) suggesting that prior loading with piRNA is necessary for the post-translational sDMA modification of Aub. We employed a Tudor-Aub binding assay that strictly depends on the Aub methylation (Kirino et al. 2010) as another tool to assess the modification status of Aub. As expected, Tudor co-immunoprecipitated with wild-type Aub, but not mdAub, from ovary lysates (Fig. 4B). Similar to mdAub, pdAub did not bind Tudor protein confirming that Aub piRNA loading is a prerequisite for sDMA modification.

**Figure 4.**
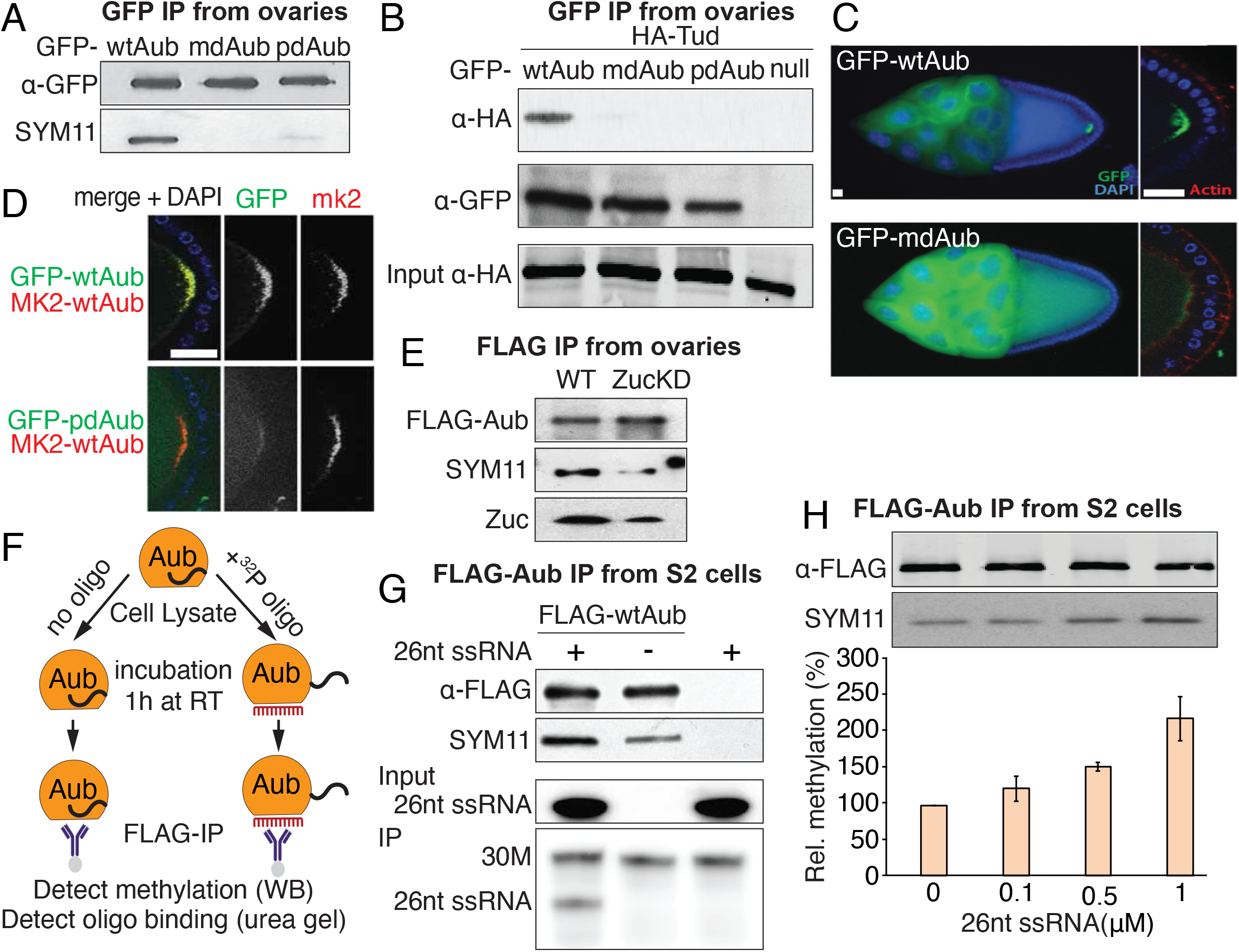
RNA binding promotes Aub arginine methylation. (A) RNA binding is required for Aub arginine methylation *in vivo*. GFP tagged wtAub, mdAub and piRNA-binding deficient pdAub were immunoprecipitated from ovaries. Transgenes were expressed on the wildtype background. Total protein level and methylation was detected in Western blot using anti-GFP and SYM11 antibodies, respectively. (B) RNA binding is required for Aub interaction with Tudor *in vivo*. HA tagged Tudor was co-expressed with GFP tagged wtAub, mdAub and pdAub. Aub was immunoprecipitated from ovarian lysate, followed by Western blot detection of the tagged transgenes. (C) Arginine methylation is required for localization of Aub to the pole plasm. GFP tagged wtAub and mdAub were separately expressed in ovaries on the wild-type Aub background. Scale bar: 5μm. (D) RNA binding is required for Aub localization into the pole plasm. Upper panel: GFP-wtAub and mKate2-wtAub were co-expressed in ovaries in the wild-type Aub background. Bottom panel: GFP-pdAub and mKate2-wtAub were co-expressed in ovaries in the wild-type Aub background. Scale bar: 5μm. (E) Zucchini KD leads to decreased Aub arginine methylation. FLAG-tagged wtAub was immunoprecipitated from control and Zuc KD ovaries in the wildtype Aub background. Methylation was detected by Western blot using the SYM11 antibody. (F) Scheme of *in vitro* oligo-binding experiments. Lysate of S2 cells expressing FLAG-Aub was incubated with or without ^32^P labelled 26nt RNA oligos followed by FLAG immunoprecipitation and detection of methylation by Western blot and oligo binding on urea gel. (G) RNA oligo binding promotes Aub sDMA modification. Aub expressed in S2 is loaded with 26nt ssRNA oligo in cell lysate, followed by immunoprecipitation. 30 nt (30M) ssRNA is spiked into each IP-ed sample to normalize for total IP-ed RNA amount. (H) Aub methylation correlates with synthetic ssRNA concentration added to lysates of S2 cells expressing FLAG-Aub. Methylation was quantified based on band intensity in FLAG and SYM11 Western blot and normalized to the no oligo control. Error bars indicate st. dev (n=2).

To further explore the link between piRNA loading and sDMA modification we tested Aub interaction with Tudor *in situ*, inside of the cells. In growing oocytes, Tudor protein localizes to the pole plasm, which is the posterior region of the oocyte essential for germ cell specification in the zygote (Arkov et al. 2006; Thomson and Lasko 2005; Santos and Lehmann 2004). Tudor binds sDMA-modified Aub leading to its accumulation in the pole plasm (Anne and Mechler 2005). As expected, while wild-type Aub was concentrated in the pole plasm, mdAub failed to be properly localized because it could not interact with Tudor protein (Fig. 4C). Similarly, to methylation-deficient Aub, pdAub had strong defects in pole plasm localization (Fig. 4D) confirming that piRNA-free Aub is unable to be methylated.

The experiments described above suggest that a defect in piRNA loading impairs Aub methylation, however it is possible that two point mutations introduced into pdAub to impair piRNA binding may cause additional structural changes that render it incapable of methylation. Therefore, we further explored the link between piRNA loading and sDMA modification using wild-type Aub protein. We analyzed the modification status of wild-type Aub in flies that have a low level of piRNAs due to knockdown of Zucchini (Zuc): a nuclease that is essential for piRNA processing (Malone et al. 2009; Han et al. 2015; Mohn, Handler, and Brennecke 2015). Depletion of Zuc in fly ovaries by RNAi caused a reduction of Aub methylation (Fig. 4E), suggesting that the modification status of wild-type Aub depends on its piRNA loading status.

To directly test if binding of guide piRNA induces Aub modification, we studied if the loading of a chemically synthesized RNA in cell lysate triggers Aub methylation. Argonaute proteins (including members of the Piwi clade) can be loaded with 5’-monophosphorylated short RNA when incubated together *in vitro* (Elkayam et al. 2012; Yamaguchi et al. 2020; Matsumoto et al. 2016). We expressed tagged Aub in S2 cells that lack an active piRNA pathway and do not express piRNA. We added different amounts of 5’-monophosphorylated 26 nt RNA into the lysate of S2 cells that expressed Aub protein and incubated lysate for one hour to allow Aub to be loaded with RNA and the methylation reaction to proceed. Next, we purified Aub and analyzed associated RNA and its modification status (Fig 4F). The labeling of bound RNA indicates that Aub binds exogenous short RNA during incubation (Fig 4G). Importantly, the addition of increasing amounts of short RNA led to a progressive increase in Aub methylation (Fig. 4H). Thus, loading of Aub with synthetic RNA triggers sDMA modification in cell lysate indicating direct link between RNA binding and modification.

### piRNA loading induces conformational change and promotes Aub modification by increased accessibility of the N-terminal sDMA motif to the PRMT5 methylosome complex

How can piRNA loading into Aub induce its methylation? Binding of RNA might induce conformational changes in protein structure that expose previously hidden residues to the solvent and enhance access to enzyme that catalyzes modification. Indeed, structural studies of Argonaute proteins demonstrated extensive conformational change in protein structure upon binding to guide RNAs (Elkayam et al. 2012; Yashiro et al. 2018). However, the structure of *Drosophila* Aub protein has yet to be solved and the N-terminal extension regions are not present in the available structures of piwi proteins (Yamaguchi et al. 2020; Matsumoto et al. 2016). To study if loading of RNA into Aub induces its conformation change, we employed the RNA loading strategy described in the previous experiment (Fig 4F) followed by a partial proteolysis assay. Aub protein with a FLAG tag introduced at its N-terminus was expressed in S2 cells and half of the lysate was incubated with 1 μM of synthetic 26 nt RNA for one hour, while RNA was omitted for the other half. After incubation, protein was captured on anti-FLAG beads and subjected to serial dilution of chymotrypsin protease followed by SDS-PAGE gel electrophoresis (Fig. S5A). Western blot with anti-FLAG antibody showed that the N-terminus of Aub has different accessibility to protease in free and RNA-bound forms (Fig S5A) suggesting that Aub undergoes a conformational change upon RNA loading.

To further explore how conformation affects Aub methylation we compared the modification status of full-length protein and N-terminal fragment that harbors the (G/A)R motif. Tagged full length Aub and the N-terminus (1-105 aa) fragment were expressed in S2 cells and the methylation status was probed by Western blot with SYM11 antibody. Similar to previous experiments, only a very weak modification was observed on full-length protein, however, we detected strong methylation of the N-terminal fragment (Fig. 5A). This result suggests that the N-terminal region of Aub protein is not easily accessible to modification if the protein is not bound to guide RNA, but the same sequence is methylated strongly if it is apart from the whole protein.

**Figure 5.**
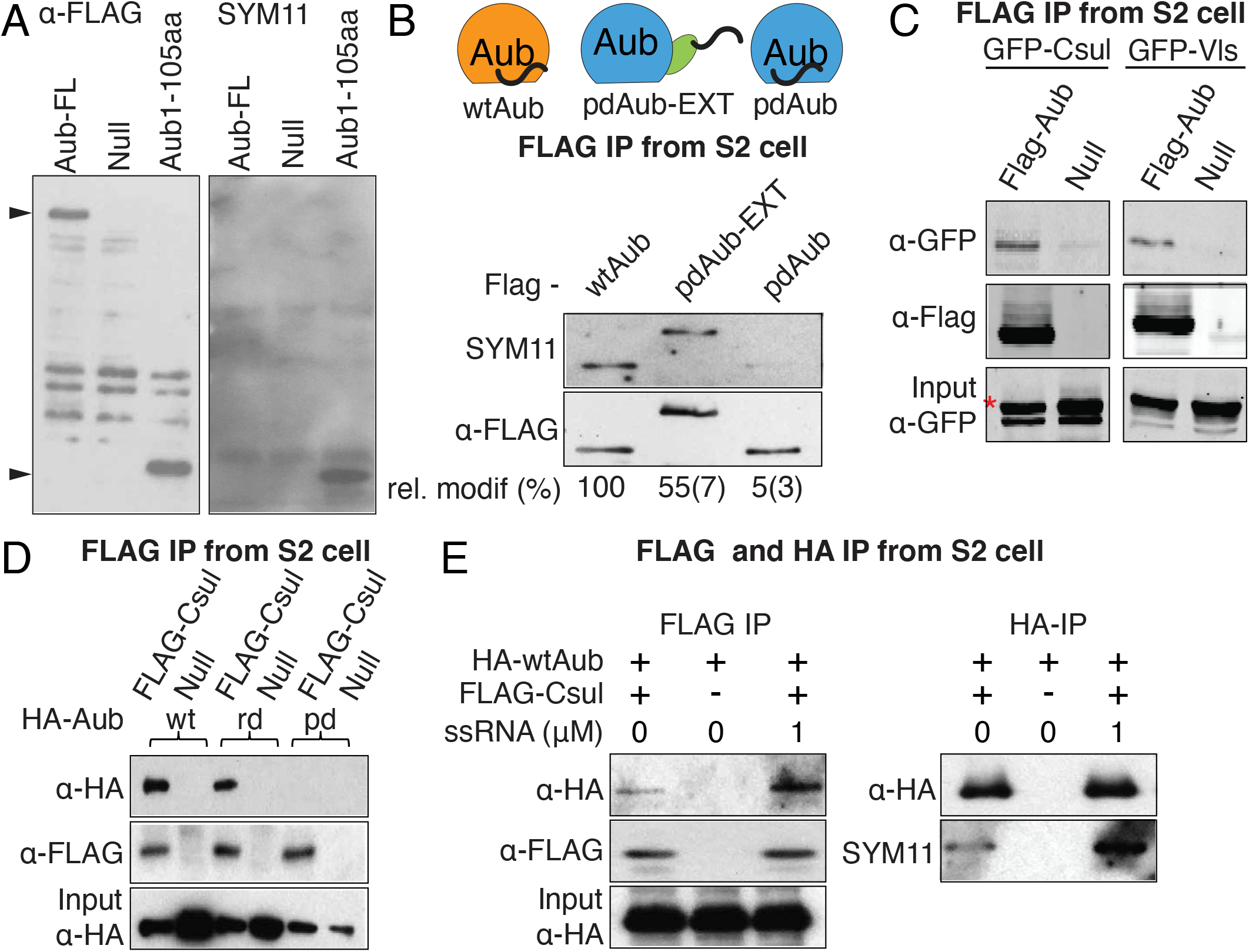
RNA binding triggers conformational change in Aub, exposing its N terminus to the methyltransferase Csul/Vls complex. (A) N-terminal region within Aub protein is not easily accessible to methylation. FLAG tagged full length and N-terminal truncated (1-105aa) Aub were expressed in S2 cell and immunoprecipitated. Total protein and methylation level was assessed by Western blot. Arrowheads indicate correct size for full-length and N-terminal fragment, respectively. (B) Top: Scheme showing architectures of different Aub constructs expressed and IP-ed from S2 cells. EGFP inserted between the N-terminal and PAZ domain (pdAub-EXT) artificially exposes N-terminus in absence of piRNA binding. Bottom: Western blot analysis of methylation states. Relative methylation level as estimated by ratio of SYM11/FLAG band intensities normalized to wildtype are listed, standard deviations are shown in brackets, n=2. (C) Aub interacts with Csul and Vls in S2 cells. Co-IP Western blot of tagged transgenes. Asterisk indicates band corresponding to GFP-Csul in the INPUT. (D) Aub binding to Csul depends on RNA binding but not on Arg methylation. FLAG-Csul and HA-Aub transgenes were expressed in S2 cells and coIP followed by Western detection. (E) RNA loading of Aub leads to increased binding to Csul and increased methylation of Aub. FLAG-Csul and HA-Aub were expressed in S2 cells, lysates were incubated in the presence or absence of ssRNA oligo prior to coIP followed by Western detection.

Next we tested if we can alter the accessibility of the N-terminal region to the methylation machinery by inserting an extra sequence between this region and the rest of the protein (named pdAub-EXT). While Aub deficient in piRNA binding was methylated only on a very weak level, insertion of a GFP sequence after AA105 caused robust methylation (Fig. 5B). Thus, insertion of an extended sequence between N-terminus and the rest of the protein converts the N-terminal (GA)R motif into a good substrate for modification even if the protein is not loaded with RNA. Overall, these experiments suggest that the N-terminal region is not readily accessible if Aub is not bound to RNA, however, it can be readily methylated upon solvent exposure.

We tested the biochemical interactions between Aub and the components of the methylosome complex responsible for sDMA modifications. sDMA modifications are catalyzed by conserved methylosome enzyme composed of *Drosophila* PRMT5 homolog Capsuleen (Csul) and MEP50 homolog Valois (Vls) (Friesen et al. 2001; Meister et al. 2001; Friesen et al. 2002) and previous genetic studies implicated both proteins are required for Aub sDMA modifications (Anne et al. 2007; Gonsalvez et al. 2006; Anne and Mechler 2005). We found that Csul and Vls co-purifies with Aub in S2 cells (Fig. 5C). We further dissected Aub protein into individual domains and found that the N-terminal fragment (1-220aa) alone was sufficient for binding to Csul, while deletion of this region disrupted the interaction (Fig. S5B). This indicated that the methylosome interacts with the N-terminal region of Aub harboring (G/A)R motif, while other regions are dispensable for binding.

To study if piRNA loading affects the interaction of Aub with the methylosome complex we tested binding of Csul with Aub mutants. Csul binds wild-type and mdAub indicating that arginine residues that are targeted for methylation are dispensable for interaction with Csul (Fig. 5D). However, Csul did not interact with the piRNA-deficient pdAub mutant (Fig. 5D) suggesting that conformation change induced by binding to RNA is required for interactions with Csul. Finally, we loaded wild-type Aub with synthesized 26 nt RNA and tested the effect of RNA binding on the interaction with Csul. Addition of synthetic RNA increased interaction of Aub with Csul and subsequent Aub methylation (Fig. 5E). Thus, we determined that increased levels of sDMA modifications upon RNA binding appears to be a direct result of stronger binding of RNA-loaded Aub to the methylosome complex.

## Discussion

Ago proteins are present in all domains of life and use a conserved molecular architecture to bind guide nucleic acids and recognize and process complementary targets (Wei et al. 2012; Song et al. 2004; Elkayam et al. 2012; Schirle and MacRae 2012). Despite the simplicity of the effector complex composed of one protein and one nucleic acid, Argonautes play crucial roles in the control of gene expression and have remarkably diverse sets of targets and functions. The diversity of post-translational modifications expands the regulation and function of Argonaute proteins. For example, phosphorylation of human Argonaute2 affects its subcellular localization and small RNA binding (Johnston et al. 2010; Zeng et al. 2008), while prolyl 4-hydroxylation is required for its stability (Qi et al. 2008). We found that a post-translational modification specific to members of the Piwi clade of the Argonaute family, sDMA in the flexible N-terminal region, encodes information about guide RNA loading status and regulates interactions, cellular localization and function of Piwi proteins.

Although Piwi proteins and piRNAs share many similarities with other Agos and their RNA guides, the piRNA pathway has evolved unique features that are essential for its function as an adaptive genome defense system. One such unique property is amplification of piRNAs that target active transposons in the ping-pong cycle (Brennecke et al. 2007; Gunawardane et al. 2007). Ping-pong employs the intrinsic RNA binding and processing capabilities of Ago proteins, however, it creates new functionality through the cooperation between two Piwi proteins. Our results indicate that the ping-pong cycle and sDMA-modification are tightly linked and that the modification status of Piwi proteins regulates assembly of the ping-pong processing complex.

### Molecular mechanism of sDMA regulation

Several lines of evidence suggest that sDMA modification of Aub is induced by binding of a piRNA guide. First, Aub that is deficient in piRNA binding has a decreased level of sDMA modification (Fig. 4A). Second, disruption of piRNA biogenesis diminishes methylation of wild-type Aub (Fig. 4E). Finally, loading of chemically synthesized RNA into Aub promotes its association with the methylosome complex (Fig. 5E) and subsequent sDMA modification (Fig. 5E). In contrast, sDMA modification of Aub is not required for its loading with piRNA (Fig. 3H,3I and S4C). Together, these results suggest that sDMA modification of Aub acts as a signal of its piRNA-bound state.

Our results suggest that piRNA loading induces sDMA methylation through a conformational change that makes the N-terminal sequence accessible to the methylation enzyme. While unloaded Aub is poorly methylated, the N-terminal sequence alone is a good substrate for methylation (Fig. 5A). Insertion of a sequence between the N-terminal region and the rest of the protein also promotes methylation (despite the protein not being able to bind piRNA), suggesting that other parts of the protein inhibit modification (Fig. 5B). Finally, partial proteolysis indicates that Aub undergoes a conformation change upon piRNA loading (Fig. S5A). Combined, these experiments suggest that the N-terminal sequence is inaccessible to the modifying enzyme until Aub binds a guide RNA, inducing a conformation change that exposes its N-terminus (Fig 6A).

**Figure 6.**
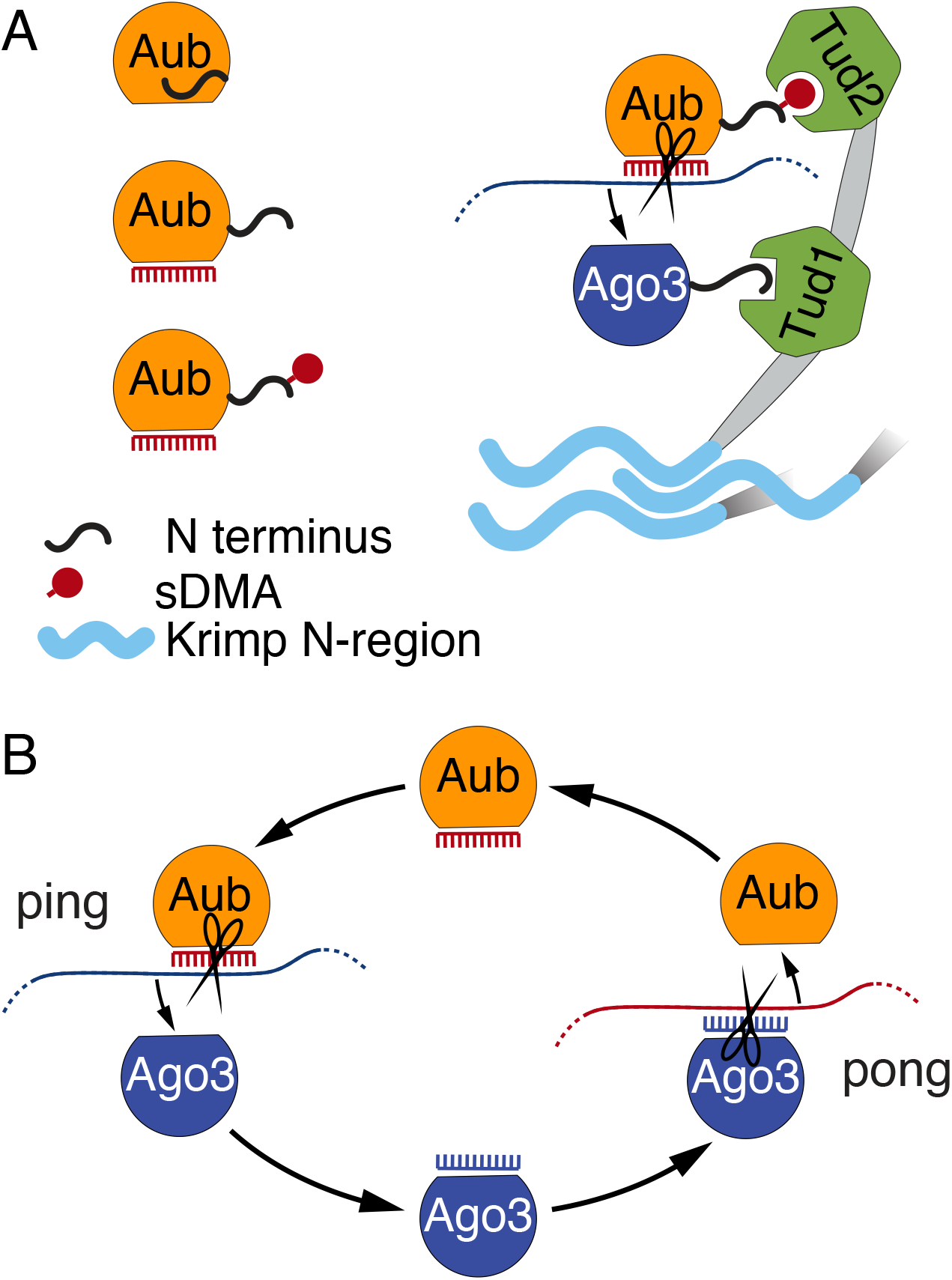
Model for sDMA regulation and its function in ping-pong cycle. (A) Model sDMA dependent assembly of the ping processing complex. The N-terminus of unloaded Aub is inaccessible. Binding to piRNA guide leads to conformational change of Aub exposing its N terminus and enabling methylation of its N-terminal arginines. Methylation thus serves as a signal of loading-state and enables Aub binding to the Tud2 domain of Krimp, which specifically recognizes methylated Arginines. The Tud1 domain of Krimp binds unmethylated, unloaded Ago3, enabling Ago3 loading with the newly processed piRNA. The N-terminal unstructured region of Krimper allows Krimper multimerization resulting in a Krimp scaffold that might assist in nuage assembly and ensuring high local concentration of Aub and Ago3. (B) The Ping-Pong cycle consists of two distinct stages. In the ping stage, Aub with its piRNA guide targets piRNA precursors or TE transcripts. Cleaved, mature piRNA is loaded into Ago3. In the pong stage, piRNA-loaded Ago3 targets antisense piRNA precursors. Cleaved mature piRNA is loaded into Aub. The ping and pong processing could be accomplished by different complexes in nuage.

Structural differences between empty and loaded states were reported for several prokaryotic and eukaryotic Agos (Elkayam et al. 2012; Rashid et al. 2007; Willkomm et al. 2017; Swarts et al. 2015; Wang et al. 2008; Miyoshi et al. 2016; Doxzen and Doudna 2017; Parker, Roe, and Barford 2005), corroborating the idea that binding to guide RNA induces conformational change. The PAZ domain of Agos exhibit high level of flexibility upon loading of guide RNA/DNA. During the recognition of target RNA, the PAZ domain undergoes a conformational transition that releases the 3’ end of the guide and facilitates downstream guide-target base pairing. Unfortunately, the N-terminal extension region was often truncated to facilitate Ago expression and crystallization and no reported structure provides information about the N-terminal extension region (Matsumoto et al. 2016; Yamaguchi et al. 2020; Nakanishi et al. 2012). If the N-terminal region is preserved, it exists in an unstructured conformation that remains unresolved by crystallography (Faehnle et al. 2013; Nakanishi et al. 2013; Schirle and MacRae 2012). However, piRNA loading of the nuclear Piwi protein in *Drosophila* was shown to induce a conformational change that exposes the nuclear localization sequence (NLS) located in its N-terminus and to enable its binding to importin (Yashiro et al. 2018). This loading-dependent accessibility of the NLS ensures that only piRNA-loaded Piwi is transported to the nucleus, where it induces chromatin repression. Thus, two Piwi clade proteins, Aub and Piwi, harbor an N-terminal sequence that becomes accessible upon piRNA loading and its exposure promotes interactions with other factors and regulates protein function. Similar to Aub, the N-terminal extension region of Ago3 also harbors a (G/A)R motif that can be modified. Considering that piRNA binding triggers exposure of the N-terminus in Aub and Piwi, a similar process might occur in Ago3. Indeed, previous studies (Sato, Iwasaki, Shibuya, et al. 2015; Webster et al. 2015) and our results revealed that, unlike the bulk of the cellular Ago3 pool, Krimp-bound Ago3 is both unloaded and unmethylated, indicating that piRNA binding and modification are correlated for Ago3 as well as for Aub.

Our results demonstrate the importance of the N-terminal region in the function of piwi proteins. Unlike other domains (PAZ, MID, PIWI) of Argonautes with well-characterized functions in RNA guide binding and target cleavage, the N-terminal region has received little attention due to its disordered conformation and its low conservation between different Agos (Meister et al. 2007; Ryazansky, Kulbachinskiy, and Aravin 2018; Wang et al. 2009; Sheng et al. 2014; Schirle and MacRae 2012; Kwak and Tomari 2012). Our results suggest that the low conservation and absence of a fixed structure are in fact important features of the N-terminal region that are critical for piwi proteins function. The flexible structure of this region might provide sensitivity to changes in overall protein conformation, such as the changes triggered by guide RNA binding. In Aub and Ago3, the modification and binding of sDMA sites to other proteins, as well as NLS-mediated interaction of Piwi require only a short linear motif, and thus the N-terminal region does not require a strongly conserved sequence or rigid folding. In agreement with this, the presence of a (G/A)R motif in Aub and Ago3 proteins is conserved in other *Drosophila* species, however, the specific position and sequence context of the motif is diverse (Fig 1D and 1E). Poor similarity between N-terminal sequences of different Agos might endow them with distinct functions. It might be worth exploring whether signaling of the guide-loading state through exposure of an N-terminal region is also conserved in Ago-clade proteins and whether it regulates their function.

### Function of sDMA in the ping-pong cycle

The central feature of ping-pong is that the cleavage of target RNA by one Piwi protein results in transfer of the cleaved product to another Piwi protein (Fig. 6B). Most eukaryotic Argonaute proteins are capable of the individual steps of this process, including RNA cleavage and binding of monophosphorylated RNA. However, there is no indication that any member of the Ago clade can engage in ping-pong, suggesting that coordination of the two steps requires additional machinery. Although the original model of ping-pong did not provide information on the molecular complex and interactions within the complex, ping-pong intuitively implies physical proximity between the two piwi proteins followed by complex molecular rearrangements. We found that information about the piRNA-loading state of piwi proteins signaled by their sDMA modifications is used to assemble a complex that enables transfer of the processed RNA from Aub to Ago3.

While our previous findings strongly suggest that Krimp plays a role in assembly of the ping-pong piRNA processing (4P) complex in which Aub and Ago3 are brought into close physical proximity (Webster et al. 2015), the architecture of this complex and the extent to which Krimp regulates ping-pong remained unknown. Our results indicate that a single Krimp molecule interacts simultaneously with Aub and Ago3, suggesting that ping-pong takes place within a tertiary complex containing one molecule of each protein. Krimp actively selects the two ping-pong partners using the distinct specificities of its two Tudor domains: eTud1 uniquely binds Ago3, while eTud2 recognizes modified Aub (Fig 1F and S1). We found that *in vitro* the eTud2 domain is capable of binding both sDMA-modified Aub and Ago3 peptides (Fig S1), however, *in vivo* Krimp complexes were reported to contain exclusively unmodified Ago3 (Sato, Iwasaki, Shibuya, et al. 2015), suggesting that in the proper cellular context eTud2 only binds sDMA-Aub. Thus, the domain architecture of Krimp ensures that tertiary complexes contain Aub-Ago3 partners rather than random pairs. This finding is in line with the observation that ping-pong occurs predominantly between Aub and Ago3 (Gunawardane et al. 2007; Brennecke et al. 2007), although, in principle ping-pong can take place between two identical proteins and a small level of homotypic Aub/Aub ping-pong was previously detected (Zhang et al. 2011; Li et al. 2009). Thus, our results suggest that the propensity for heterotypic ping-pong is, at least in part, due to Krimp (Fig. 6A).

Ping-pong not only requires the physical proximity of two piwi proteins but also that they have opposite piRNA-loading states: one protein induces piRNA-guided RNA cleavage (and therefore has to be loaded with a piRNA guide), while the other accepts the product of this reaction (and therefore has to be free of piRNA). Our results suggest that the opposite binding preference of the two Tudor domains towards sDMA ensures that the tertiary complex contains piwi proteins in opposite RNA-loading states. While the overall fold structure of the two Tudor domains is similar, they have critical differences responsible for their distinct binding preferences. The binding pocket of eTud2 is similar to that of other Tudor domains and contains four aromatic residues that interact with sDMA (Fig. 2A and 2D). As sDMA modification of Aub signals its piRNA-binding status, the binding of eTud2 to modified Aub ensures that the complex contains Aub/piRNA. The structural studies and *in vitro* binding assays revealed that Ago3 binds to eTud1 in its unmethylated state and sDMA modification of any of the Arg residues within its (A/G)R motif prevents this interaction. The unusual binding preference of eTud1 is reflected in its non-canonical binding pocket, which lacks three of the four conserved aromatic residues (Fig. 2H). Binding of methylated Aub and unmethylated Ago3 ensures that Aub has guide piRNA and Ago3 is free, thus enabling loading of Ago3 with RNA generated by Aub/piRNA-induced cleavage (Fig. 6A).

The architecture of the tertiary complex assembled by Krimp permits Aub-dependent generation and loading of RNA into Ago3. However, the ping-pong cycle also includes the opposite step, Ago3-dependent generation of Aub piRNA (henceforth we termed these steps ‘ping’ and ‘pong’). The majority of Ago3-bound piRNA (produced by ping) is in sense orientation relative to transposon mRNAs and therefore cannot directly repress these transcripts (Brennecke et al. 2007). It is the latter ‘pong’ step of ping-pong that generates new antisense piRNAs responsible for targeting transposons and therefore endows the cycle with its biological function. This pong step also requires Aub-Ago3 interaction, however, piRNA-binding and therefore the methylation statuses of the two proteins must be reversed. How is this pong complex assembled? One possibility is that a different protein binds Aub and Ago3 in their opposite modification states. Genetic studies identified several Tudor-domain proteins involved in piRNA biogenesis and some of them, such as Qin and Tapas (Zhang et al. 2011; Patil et al. 2014), were shown to be required for the ping-pong cycle. It will be interesting to see if one of these proteins assembles the pong complex. Our results suggest that the ping and pong steps require assembly of two distinct complexes discriminated by the modification status of Aub and Ago3.

### Formation of a membraneless cellular compartment and its function in ping-pong

As a single Krimp simultaneously binds Aub and Ago3, Krimp dimerization might be dispensable for ping-pong, raising the question what the function of Krimp self-interaction is. Previous findings suggest that Krimp forms a scaffold for assembly of nuage, a membraneless organelle (MLO) that surrounds nuclei of nurse cells and resembles other MLO possibly formed through liquid-liquid phase separation (Webster et al. 2015). Several lines of evidence point at Krimp as an essential component of nuage that acts as a scaffold for its assembly and the recruitment of client components. First, unlike other nuage components, FRAP measurements show very little Krimp exchange between nuage and the dispersed cytoplasmic compartment. Second, wild type, but not mutant Krimp that lacks the self-interaction domain, forms cytoplasmic granules upon expression in heterologous cells that do not contain other nuage proteins. In contrast, other nuage components including Aub and Ago3 are dispersed in the cytoplasm when expressed in a similar setting, suggesting that they do not form condensates on their own and rely on other components for recruitment to nuage. Krimp recruits both Aub and Ago3 into MLO that it forms in heterologous cells. Combined, these data indicating that Krimp works as a scaffold and Ago3 and Aub as its clients for nuage assembly. Thus, the interactions between Krimp and the N-terminal regions of Aub and Ago3 is not only essential for the assembly of the tertiary molecular complex but is also responsible for recruitment of these proteins into membraneless cellular compartment (Fig. 6A). The high local concentration of proteins and RNA involved in the piRNA pathway in nuage might enhance the efficiency of ping-pong as well as recognition of RNA targets by Aub and Ago3.

## Acknowledgements

We thank members of the Aravin lab for discussion and comments. We thank the BL19U1 beamlines staff at the Shanghai Synchrotron Radiation Facility for assistance during data collection. We thank Igor Antoshechkin (Caltech) for help with sequencing. This work was supported by grants from the National Institutes of Health (R01 GM097363) and by the HHMI Faculty Scholar Award to A.A.A. and the National Natural Science Foundation of China (31870755) and the Guangdong Innovation Research Team Fund (2016ZT06S172) to S.L.

## Materials and Methods

### Fly stocks

Short hairpin RNA (shRNA) lines used for knockdown including sh-White (BDSC #33623) and sh-Zuc (BDSC #35227), maternal alpha-Tubulin 67C-Gal4 drivers on chromosome two (BDSC #7062) or chromosome three (BDSC #7063) in addition to the Aub mutant stocks aub^HN2^ cn^1^ bw^1^/CyO (BDSC #8517) and aub^QC42^ cn^1^ bw^1^/CyO, (BDSC # 4968) were obtained from the Bloomington *Drosophila* Stock Center. Flies were kept on yeast for 2 days and ovaries dissected 5 days after hatching.

### Generation of Transgenic Fly Lines

Transgenic constructs for injection were generated using the Gateway cloning system (Life Technologies). cDNAs were obtained by RT-PCR from ovarian or testes RNA of adult *Drosophila melanogaster,* Oregon R strain. mdAub and pdAub were generated by overlap PCR and inserted into the pENTR-D-TOPO directional cloning vector (Life Technologies). Transgenes were cloned into the pUASP-Gateway-phiC31 fly injection vector derived from the pCasPeR5-phiC31 vector containing GFP, mKate2 or Strep-FLAG tags using the Gateway cloning system (Life Technologies). The expression of each transgene was controlled using the yeast upstream activation sequence promoter (UASp) stably crossed with a maternal a-Tubulin67c-Gal4-VP16 (MaG4) driver. Transgenes were generated in flies by PhiC31-mediated transformation (BestGene) using PhiC31 landing pads on either chromosome two (BDSC #9736) or chromosome three (BDSC #9750). The GFP-wtAub and GFP-mdAub BAC line was generated by cloning of the *aub* genomic locus from the BAC clone BACN04M10 into the pCasPeR4 vector using restriction sites XhoI and SpeI. Bacterial recombineering (Gene Bridges Counter Selection kit) was used to insert an in-frame GFP tag in the start site of Aub. GFP-wtAub and GFP-mdAub rescue lines were generated by crossing transgenic construct into the aub^[HN]^/ aub^[QC]^ background.

### Cell Culture, Immunoprecipitation and Western Blots

Schneider S2 cells were cultured in complete Schneider medium (10% heat inactivated FBS; 100U penicillin [Life technologies]; 100μg streptomycin [Life technologies]). Plasmids were generated using Gateway cloning (Life technologies) using the *Drosophila* Gateway Vector Collection (DGVC) destination vectors, pAGW, pAFW and pAHW for GFP, 3xFLAG and 3XHA tags, respectively, driven by the Actin5C promoter. All constructs were N terminal tagged. Cells were transfected using TransIT-LT1 transfection reagent (Mirus biosciences) according to the manufacturer’s recommendation using 3μg of total plasmid, with 1.5μg each for double transfections, or 1.0μg each for triple transfections. S2 cells were lysed in S2 lysis buffer (20mM Tris at pH7.4, 200mM KCl, 0.1% Tween-20,0.1% Igepal, EDTA-free Complete Protease Inhibitor Cocktail [Roche], 100μg/mL RNase A). Supernatant was cleared by centrifugation at 4,000 x g for 20 minutes at 4°C. Input sample was collected from the supernatant at concentrations of 1-3μg/μL. Anti-FLAG M2 beads (Sigma Aldrich), anti-HA agarose beads (Thermo Fisher) and anti-GFP antibody (Covance) conjugated to Dynabeads (Thermo Fisher) were blocked in 5mg/ml BSA for 10 minutes at 4°C, followed by washing in S2 lysis buffer. Beads were added to the supernatant and rotated at 4°C for 4h, washed three times in lysis buffer and eluted by boiling in reducing SDS loading buffer. In the two-step co-IP, FLAG beads were eluted using 3X FLAF peptide (Sigma-Aldrich).

For co-IP from ovaries, dissected ovaries were lysed in NT2 buffer (50 mM Tris at pH 7.4, 150 mM NaCl, 1mM MgCl_2_, 0.05% Igepal, EDTA-free Complete Protease Inhibitor Cocktail), while for IP experiments RIPA buffer (25 mM Tris at pH 7.4, 150 mM NaCl, 1% NP-40, 0.5 sodium deoxycholate and 0.1% SDS, Igepal, EDTA-free Complete Protease Inhibitor Cocktail) was used for lysis. Lysate was incubated in the presence or absence of 100μg/mL RNase A and cleared by centrifugation. For co-IP experiments, lysates were incubated with anti-FLAG M2 beads (Sigma Aldrich) or with anti-GFP antibody (Covance) conjugated to Dynabeads (Thermo Fisher) at 4°C for 4h. For IP experiments, lysates were incubated with GFP nanobody (Chromo Tek), followed by washing and elution in reducing SDS buffer. Western blot were probed with rabbit anti-GFP (Covance) (1:3K), rabbit anti-Zuc (Endogenous) (1:1K), rabbit SYM11 antibody (Sigma Aldrich) (1:1K), mouse anti-GFP (Santa Cruz) (1:3K), anti-FLAG M2 (Sigma Aldrich) (1:10K), mouse anti-HA (Sigma Aldrich) (1:10K).

### piRNA isolation from immunopurified protein complexes

Immunopurified protein-RNA complexes were spiked with 5pmol of 42-nt RNA oligonucleotide, followed by proteinase K digestion and phenol extraction. Isolated RNA was CIP treated, radiolabeled using PNK and gamma-P32 ATP, and run on a 15% urea-PAGE gel. Semi-Quantitative piRNA binding analysis and a detailed protocol can be found in a previous paper (Webster et al. 2015).

### Small RNA-seq

Small RNA libraries were cloned from total ovarian lysate of Aub heterozygous, aub^[HN]^/ aub^[QC]^ mutant and aub^[HN]^/ aub^[QC]^, GFP-wtAub or GFP-mdAub rescue flies. 30 ovaries were dissected, and total RNA was isolated with Ribozol (Amresco, N580). Small RNAs within a 19-to 29-nt window were isolated from 15% polyacrylamide gels from 4 μg of ovarian total RNA. For samples that were NaIO_4_-treated, 5× borate buffer (pH 8.6; 150 mM borax, 150 mM boric acid) and 200 mM sodium periodate were added to the size-selected small RNA, and the samples were incubated for 30 min at 25°C. The NaIO_4_-treated small RNA was then ethanol-precipitated before proceeding to library construction. The small RNA libraries were constructed using the NEBNext small RNA library preparation set for Illumina (no. E7330S), according to the protocol, using NEBNext multiplex oligos for Illumina (no. E7335S). Libraries were sequenced on the Illumina HiSeq 2500 (SE 50-bp reads) platform.

### Immunoprecipitation small RNA-seq

Ovaries (∼100 per immunoprecipitation) from flies expressing GFP-wtAub and GFP-mdAub under the control of endogenous promoter were dissected and lysed on ice in 250 μL of lysis buffer [30 mM Hepes-KOH at pH7.4, 100 mM KOAc, 2 mM Mg(OAc)_2_, 5 mM DTT, 0.5% (v/v) NP40, proteinase inhibitor (Roche, 11836170001), RNasin Plus (Promega, N2611)]. Lysate was dounced and clarified by centrifugation at maximum speed at 4°C. The supernatant was incubated with rabbit polyclonal anti-GFP (Covance) conjugated to Dynabeads (Thermo Fisher) for 4 h at 4°C. The immunoprecipitation and RNA isolation were carried out as described previously (Vagin et al. 2006). A fifth of the RNA was CIP-treated (New England Biolabs, M0290S) in NEB buffer #3 (New England Biolabs, B7003S) for 30 min at 37°C and then ethanol-precipitated after phenol:chloroform and chloroform extraction. The CIP-treated RNA was then PNK-treated with 1 μL of 10× T4 polynucleotide kinase buffer (New England Biolabs, B0201S), 2 μL of [γ-P^32^]ATP (PerkinElmer, BLU502A250UC), and 1 μL of T4 polynucleotide kinase (New England Biolabs, M0201S) for 45 min at 37°C. The CIP- and PNK-treated RNA was added back to the remainder of the RNA isolated from the immunoprecipitation. Size selection, library preparation, and analysis were performed as described in small RNA-seq, except that fragments were gel-extracted based on labeled immunoprecipitation material.

### Sequence analysis of piRNA libraries

Sequenced total small RNA libraries from Aub heterozygous control, wtAub and mdAub rescue as well as aub^[HN]^/ aub^[QC]^ mutant flies were filtered for reads 23-29nt in length and mapped to annotated repeat sequences. TOP20 TE families with most abundant reads in the control ovaries were used for the further analysis. The fold-change in read count for each TE family was calculated by obtaining the base 2 logarithm ratio of reads in experimental libraries and heterozygous control libraries. Read counts corresponding to sense piRNA were obtained by filtering repeat sequences 23-29 nt in length for reads that were sense orientation to annotated repeats and contained adenine (A) at position 10 and not uridine (U) at position 1. The ping-pong pair fraction was obtained by the same analysis as previous study (Webster et al. 2015). Unique reads corresponding to annotated TE families were mapped to piRNA clusters 42AB, 38C, 80EF and Flam. Sequenced immunoprecipitated small RNA libraries from ovaries expressing GFP-wtAub and GFP-mdAub in wild-type background were filtered for reads 23-29 in length and mapped to annotated repeat sequences. TOP20 TE families with most abundant reads in GFP-wtAub ovaries were used for antisense ratio analysis.

### Microscopy

Ovaries were dissected in PBS and fixed in 4% PFA in PBS for 20 minutes, washed 3×10 min at RT, permeabilized in 1% Triton-X100 in PBS for 10 minutes, and DAPI stained (Sigma-Aldrich). Ovaries were washed in PBS and mounted in Vectashield medium (Vector Labs). S2 cells were allowed to settle on coverslips treated with Poly-L-Lysine (Sigma-Aldrich). After gentle washing, cells were fixed in 0.5% PFA in PBS for 20 minutes followed by staining with DAPI (Sigma-Aldrich), washed and mounted in Vectashield medium (VectorLabs). Images were captured using an AxioImager microscope; an Apotome structured illumination system was used for optical sections (Carl Zeiss).

### Limited protease assay

10ng/ul chymotrypsin stock solution was prepared in chymotrypsin reaction buffer (10 mM Tris-HCl [pH 8.0], 2 mM CaCl_2_, and 5% glycerol). FLAG-wtAub was expressed in S2 cells. Lysates were divided into two equal fractions, one was incubated with 26nt ssRNA oligo for 1h at RT, another was incubated without oligos, followed by immunoprecipitation using anti-FLAG M2 beads. Beads were washed three times with lysis buffer and then incubated with a 1:2K, 1:4K and 1:8K serial dilution of the thermolysin protease for 30 min at 37°C. After extensive washing, beads were eluted in reducing SDS buffer. Samples were analyzed by western blot using anti-FLAF M2 antibody.

### Protein disorder prediction and conservation analysis

Disorder predictions of full-length Krimper and the N terminal region of Aub and Ago3 were obtained using the PrDOS server based on a previously used algorithm (Ishida and Kinoshita 2007). Conservation is measured as a numerical index reflecting the conservation of physico-chemical properties of amino acids in the alignment: Identities score highest, followed by substitutions to amino acids lying in the same physico-chemical class. A detailed description of the algorithm can be found in (Livingstone and Barton 1993).

### Hatching rate calculation

3-day old mated adult female flies fed on yeast paste were transferred to fresh grape agar plates and allowed to lay eggs for 12 hours. Eggs were counted, and the hatching rate was determined over the following 36h hours. Counting was repeated on 10 consecutive days.

### RT-qPCR

Total RNA was isolated from 20 ovaries with Ribozol (Amresco, N580). RNA was DNase I treated (Invitrogen, 18068-015) and reverse transcribed using SuperScript III (Invitrogen) with oligo d(T)_15_ according to the manufacturer’s recommendations. RT-qPCR was performed using Mastercycler ep Realplex PCR thermal cycler machine (Eppendorf). Target expression was normalized to *rp49*. Primers are shown in Table S2.

### Fluorescence Recovery After Photobleaching (FRAP)

For each construct, at least 20 independent FRAP experiments were performed using ovaries expressing a single GFP-tagged transgene under the endogenous promoter. FRAP experiments were captured on a Zeiss LSM710 confocal microscope (Carl Zeiss AIM) equipped with a 25x/0.8 NA Imm. Corr. Multi-immersion objective lens using the Zeiss Zen Black software. Image acquisition for all experiments utilized an identical 488nm AOTF laser power setting of 7% to ensure laser power was not influencing measurements. PMT gain settings were variably set to accommodate the expression level of each GFP tagged protein. Images were acquired at 256 × 256-pixel resolution at 0.07μm pixel size and scan speed of 614.4 ms per frame with 1.0 μs pixel dwell time. A single bleach region was defined for each experiment, consisting of a region of 7×7 pixels equal to 0.49 μm x 0.49 μm and was bleached by a single iteration of 100% laser power from 488, 561 and 633nm wavelengths. Five initial pre-bleach images were captured prior to bleaching and 115 subsequent postbleach images were acquired every 614.4ms to assess fluorescence recovery in the bleach zone. FIJI (FIJI; http://fiji.sc/) software was used to analyze FRAP experiments. Detailed analysis of the mobile fraction and the nuage/cytoplasm ratio can be found in a previous study (Webster et al. 2015).

### Oligo binding in cell lysates and measurement of methylation level

S2 cells were transfected with 10μg of plasmid expressing N terminally tagged FLAG-Aub driven by the Actin5C promoter. After 48h incubation, cells were lysed in S2 lysis buffer (20mM Tris at pH7.4, 200mM KCl, 0.1% Tween-20,0.1% Igepal, EDTA-free Complete Protease Inhibitor Cocktail [Roche], 100μg/mL RNase A). Lysates were divided into four equal fractions. 26nt synthetic RNA oligo (IDT) labelled with [γ-P^32^]ATP (PerkinElmer, BLU502A250UC) as described above was added into lysate fractions to final concentration 1μM. Lysates were Incubated at RT for 1h followed by IP with FLAG M2 beads at 4°C for 4h. Half of the beads were subject to RNA isolation, the other half were used for western blot. In the oligo binding concentration gradient experiment, lysates were divided into four equal fractions. non-radioactivity labelled 5’-end phosphorylated ssRNA oligo (IDT) was added into each fraction to a final concentration of 0, 0.1, 0.5 and 1μM, respectively. Lysates were incubated at RT for 1h followed by IP at 4°C for 4h. Protein and methylation levels were detected by Western blot and methylation level estimated using FIJI based on the intensity of the protein band I_aub_ and background I_b1_ as well as the intensity of methylation signal I_m_ and background I_b2_:

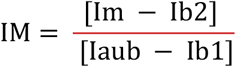

Relative methylation level was measured by normalizing to the methylation level of the no oligo control.

### Protein expression and purification

Krimper eTud1 (residues 272-512) was cloned into a self-modified pSumo vector with 10xHis tag followed by a yeast sumo sequence. The plasmid was transformed into *E. coli* strain BL21(DE3) Rosseta and cultured at 37 °C in LB medium. The protein expression was induced by adding IPTG to a final concentration of 0.2 mM when the OD600 reached 0.7, and the cells were cooled to 16 °C. The recombinant expressed protein was purified using a HisTrap column (GE Healthcare). The hexahistidine plus yeast sumo tag was removed by ulp1 protease digestion followed by a second step HisTrap column (GE Healthcare). The target protein was further purified using MonoQ and Superdex G75 columns (GE Healthcare).

Krimper eTud2 (residues 562-746) was cloned into a self-modified His-MBP vector. Protein production procedure was to with Krimper eTud1. The recombinant expressed protein was purified using a HisTrap column (GE Healthcare). The hexahistidine plus MBP tag was removed by TEV protease digestion followed by a second step HisTrap column (GE Healthcare). Protein was further purified using Q and Superdex G75 columns (GE Healthcare).

### Crystallization and crystal growth

Krimper eTud1 (272-512) was concentrated to 20 mg/ml and screened in sitting drop at 4 °C. Crystals were grown for 5 days in 0.1 M HEPES pH 7.5, 1.26 M (NH4)_2_SO_4_. For obtaining the protein-peptide complex, 20 mg/ml Krimper eTud1 was mixed with Ago3-2 peptide with a molar ratio of 1:4 and incubated for 1h before setting up sitting drop screening at 4 °C. Crystals were grown for 5 days in 0.1 M HEPES pH 7.5, 1.5 Li_2_SO_4_. 10 mg/ml Krimper eTud2 was mixed with AubR15me2 peptide with a molar ratio of 1:4 and incubated for 1h before sitting drop. Crystals appeared in 2 days in 16% PEG3350 and 0.1 M NH4Ac.

### Data collection and structure determination

All data were collected at the Shanghai Synchrotron Radiation Facilities beamline 18U1 (SSRF-BL18U1) and beamline 19U1 (SSRF-BL19U1). Data were processed using the program HKL3000 (Minor et al. 2006). Structures of eTud1-Ago3 complex and eTud2-Aub complex were solved by SAD using the anomalous signals of SeMet using the program Phenix (Adams et al. 2010). eTud1 apo structure was solved by molecular replacement using the program Phenix (Adams et al. 2010) with eTud1-Ago3 complex as a search model. Models were refined in Phenix (Adams et al. 2010) and COOT (Emsley et al. 2010) iteratively and finally presented using Pymol (DeLano Scientific).

### ITC

All ITC was performed using the Malvern PEAQ ITC instrument (Malvern) in a buffer of 50 mM NaCl, 20 mM HEPES pH 7.5 and 2 mM β-mercaptoethanol. Data analysis was performed using the Malvern data analysis software and Origin 7.0.

#### Data availability

Small RNA-seq data are available on the GEO database, GSE153156.

X-ray structures have been deposited in the RCSB Protein Data Bank with the accession codes: 7CFB for the eTud1 apo structure, 7CFC for the eTud1-Ago3 complex structure and 7CFD for the eTud2-AubR15me2 structure.

## Supplementary Figures

**Figure S1.**
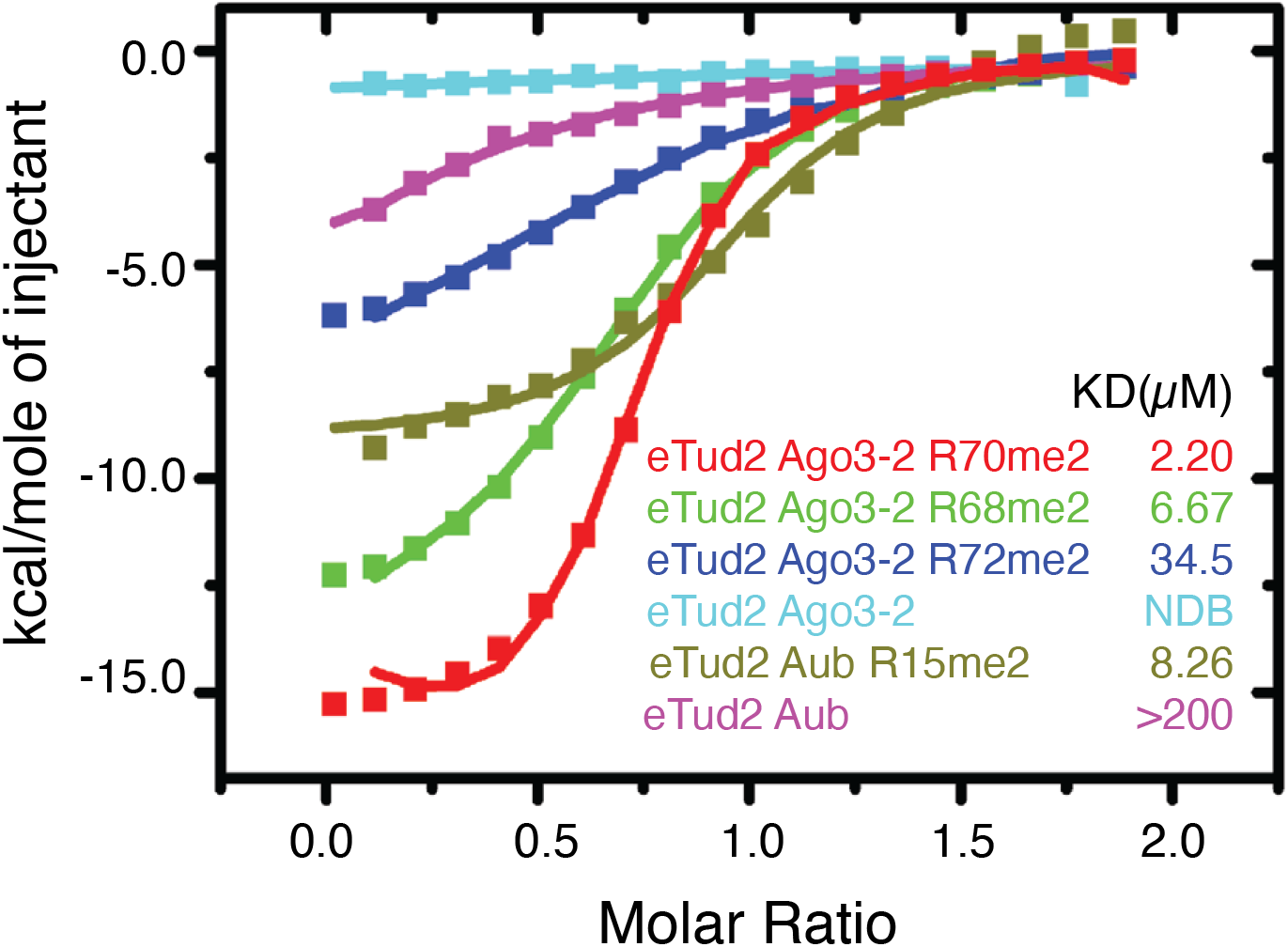
eTud2 preferentially binds methylated peptides. (A) ITC analysis of Krimper eTud2 interaction with different Ago3 or Aub peptides.

**Figure S2.**
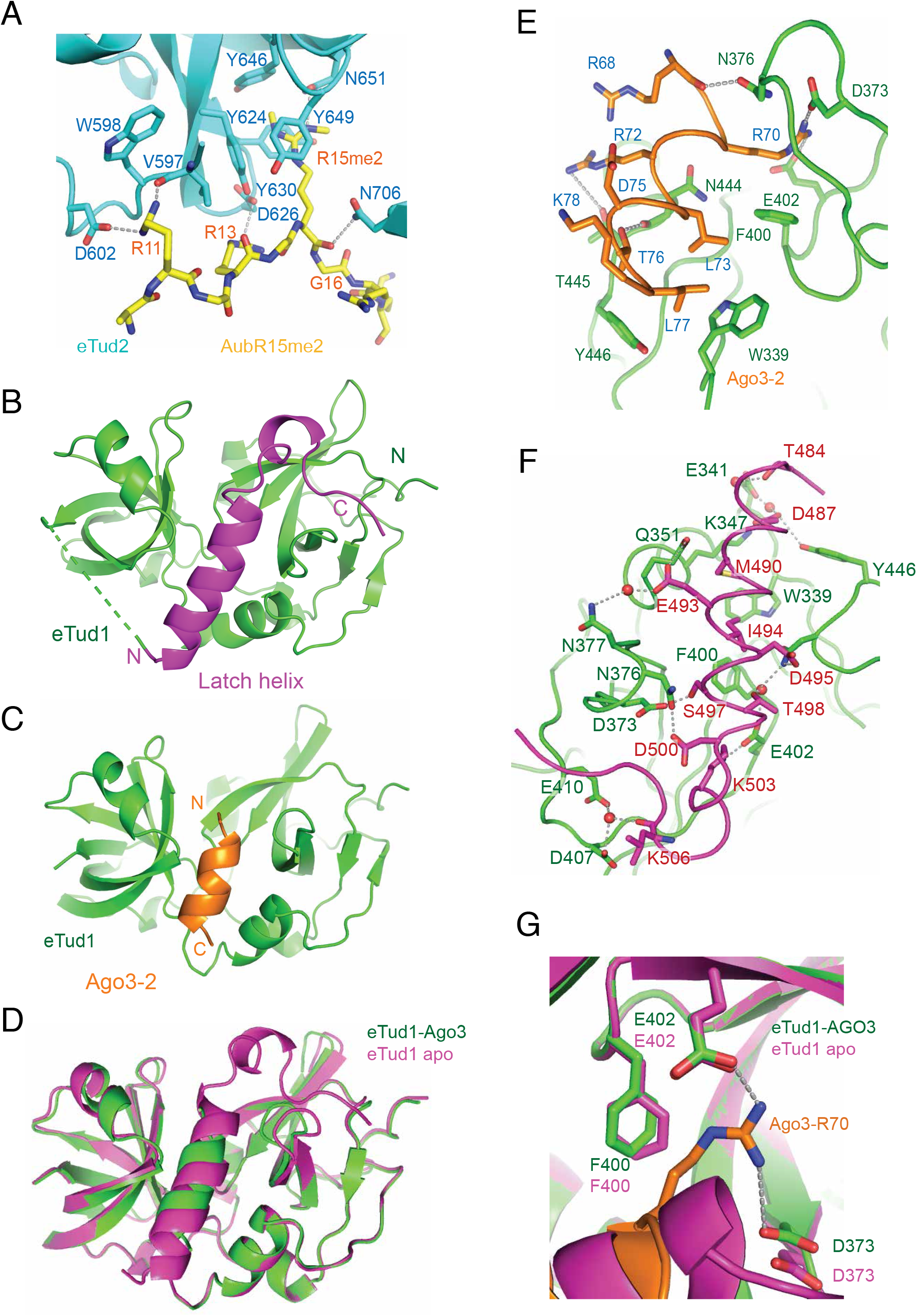
Detailed analysis of Krimper eTud1 apo structure. (A) Detailed interactions of eTud2 with Aub peptide. Hydrogen bondings are shown in grey dashed lines. (B) Overall structure of the Krimper eTud1 domain. The ‘latch helix’ region is highlighted in magenta. (C) Overall structure of the Krimper eTud1-Ago3-2 complex with Ago3-2 shown in orange. (D) Structural superposition of Krimper eTud1 apo and Ago3 peptide-bound structure. (E) Detailed interaction networks of Krimper eTud1 with the Ago3-2 peptide. (F) Detailed interactions of the ‘latch helix’ with other parts of Krimper eTud1. (G) The Phe400 side chain in the Ago3 peptide bound structure (green) rotates 90° to accommodate Ago3-R70 when compared with the apo structure (magenta).

**Figure S3.**
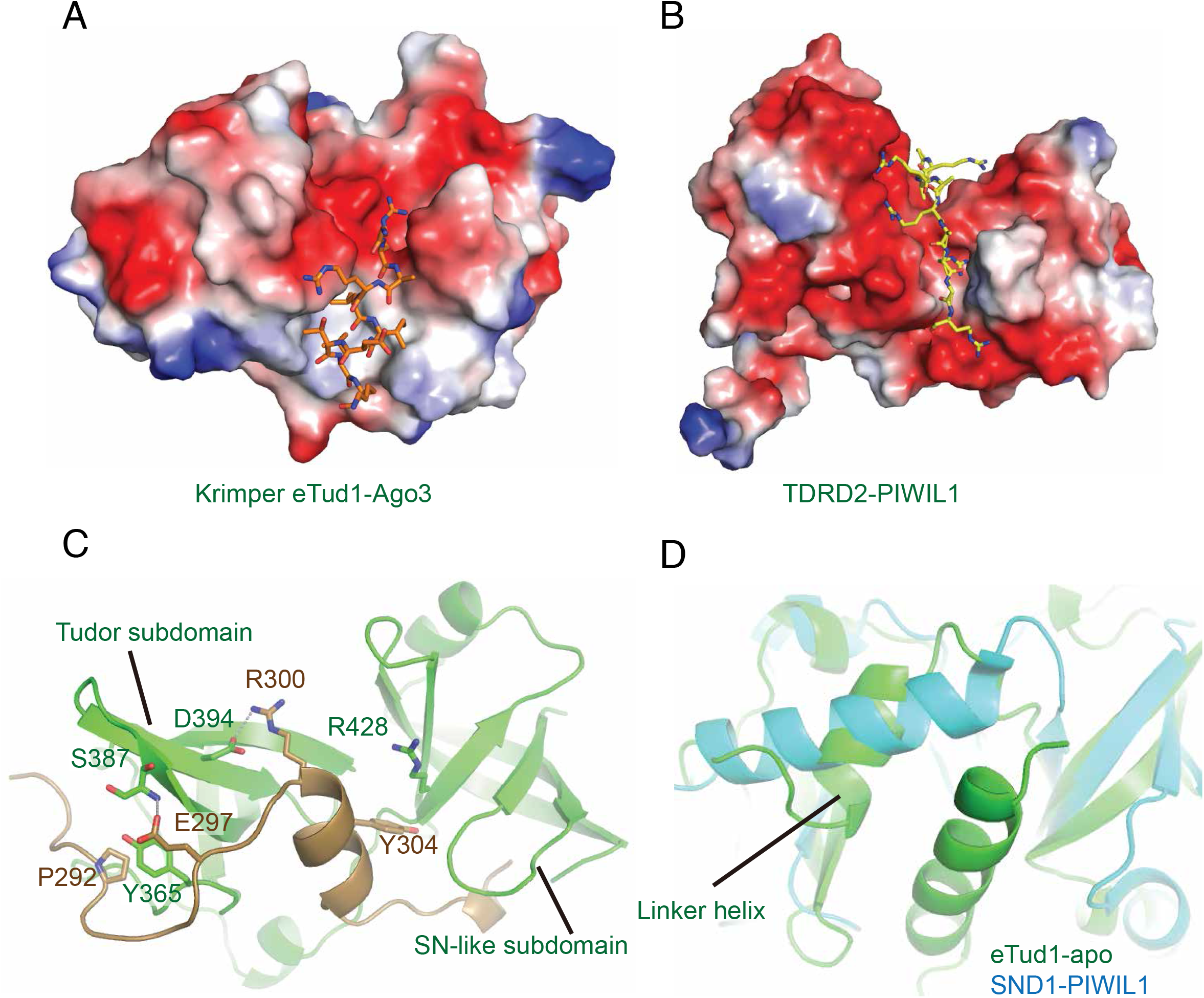
Structural comparison of Krimper eTud1-Ago3-2 complex with other extended tudor domains. (A) Electrostatic surface of the eTud1-Ago3 complex binding cleft (B) Electrostatic surface of the TDRD2-PIWIL1 complex binding cleft (PDB code:6B57) (C) eTud1 N-terminal structure is highlighted in brown. (D) Structural superposition of eTud1 apo structure with a canonical extended Tudor domain (SND1-PIWIL1, PDB code: 3OMC). The position of the linker helix is different in the two structures.

**Figure S4.**
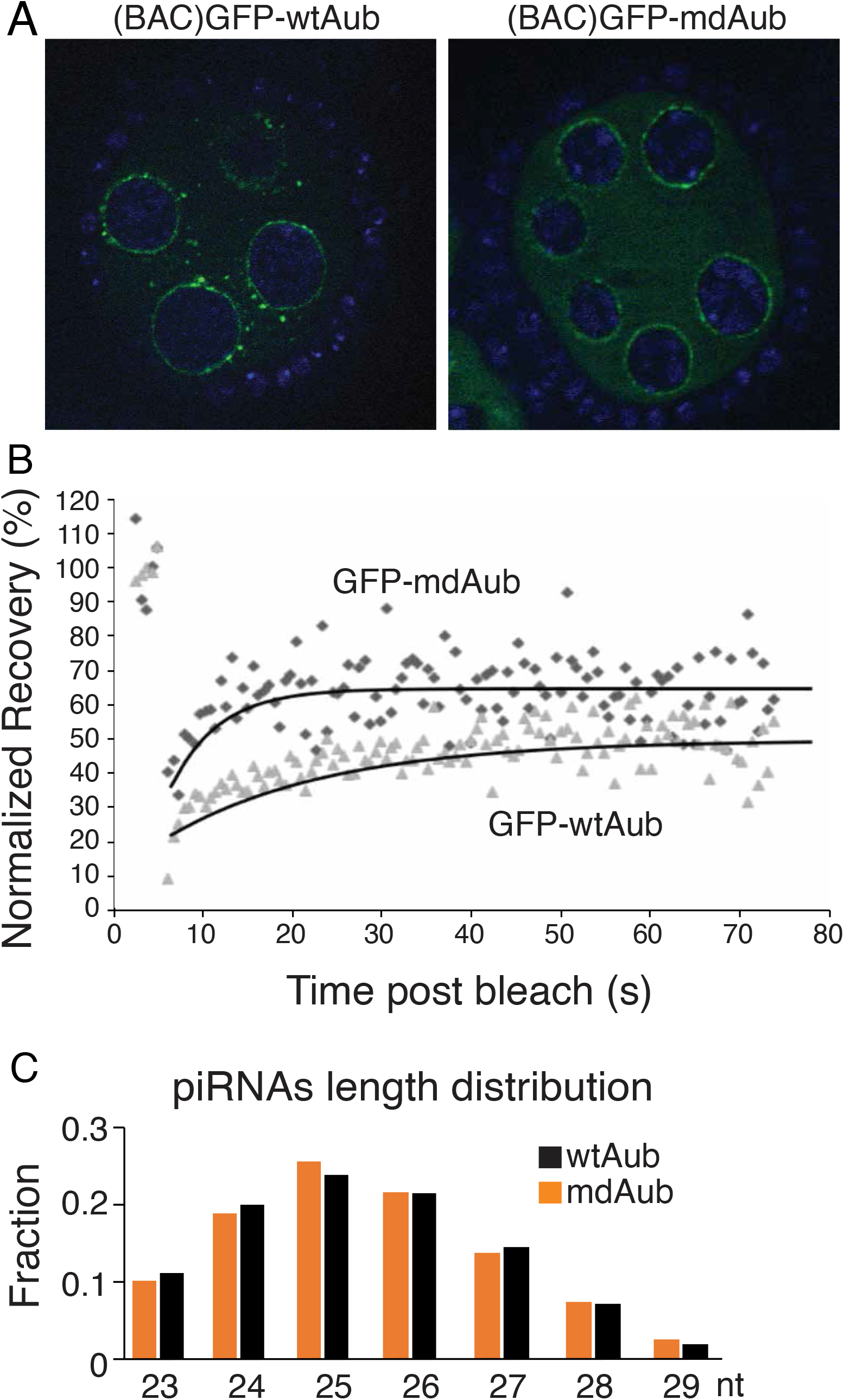
Methylation deficiency increases Aub mobile fraction. (A) Localization of GFP-wtAub and GFP-mdAub expressed under the control of the endogenous Aub promoter in nurse cells at stage 6 of oogenesis. (B) Representative FRAP experiments showing that the normalized recovery of GFP-mdAub is approximately 10% greater compared to GFP-wtAub. The mobile fraction was determined by modeling the recovery to an exponential recovery curve. (C) Length distribution of small RNAs bound by GFP-wtAub and GFP-mdAub in the wildtype Aub background.

**Figure S5.**
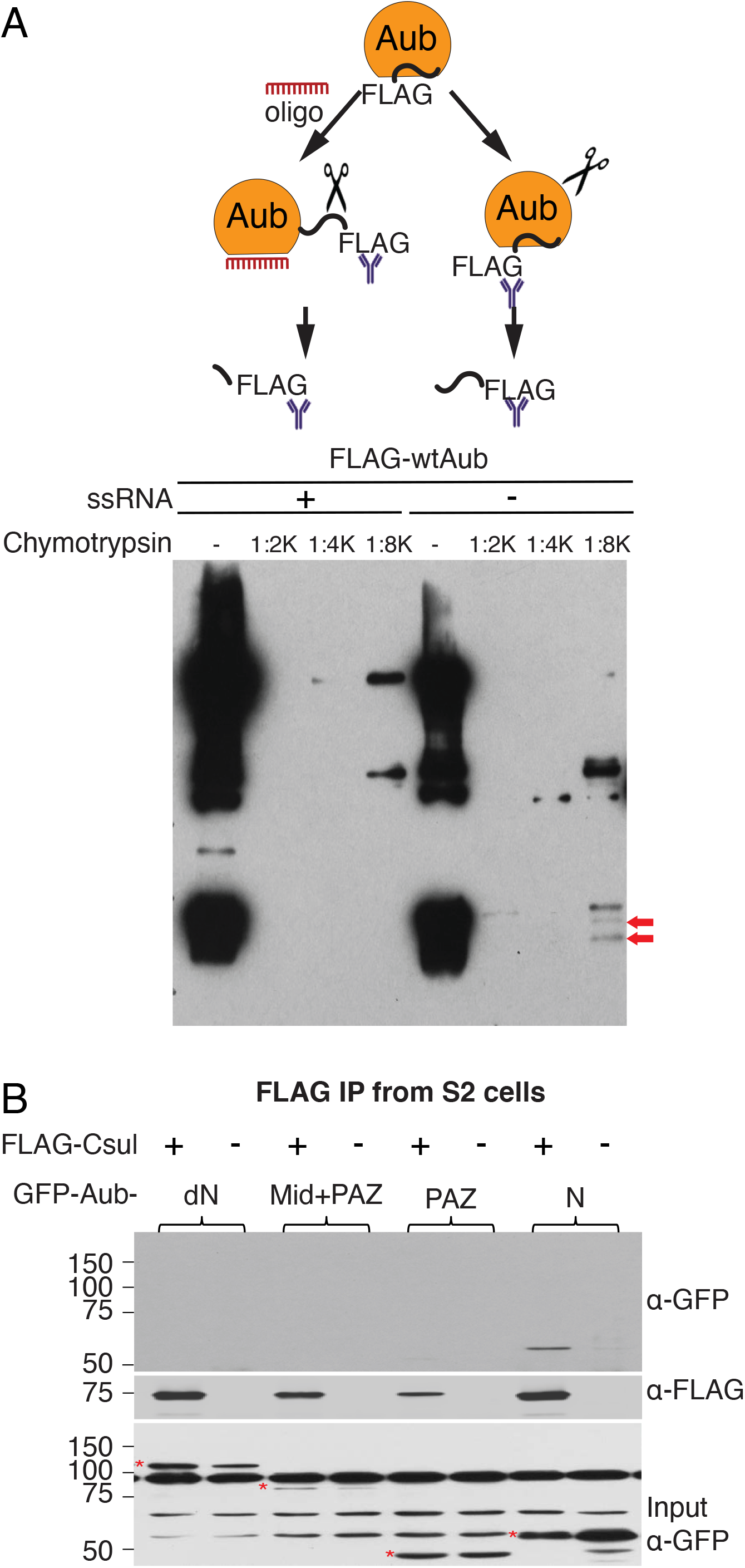
Oligo binding causes Aub conformational change. (A) Scheme of limited protease assay (top). S2 cell lysate expressing FLAG-Aub was incubated with or without 1μM synthetic ssRNA oligos followed by FLAG IP. Bead-bound proteins were incubated with different concentration of chymotrypsin. Bead fraction was analyzed by Western blot (bottom) using FLAG detection. Red arrow indicates FLAG-tagged N-terminal fragment that is undigested in the absence of piRNA loading, but digested upon small RNA loading. (B) Csul interacts with the N-terminus of Aub. FLAG tagged Csul was co-expressed in S2 cells with different GFP-tagged Aub fragments and coIP followed by Western detection of IP-ed proteins. Asterisk indicates bands corresponding to indicated GFP-tagged Aub protein fragments.

## Supplementary Tables

**Table S1. Structural Data collection and refinement statistics**

**Table S2. Oligo and primer sequences**

---

**Table 1.** Data collection and refinement statistics

|  | Krimper Tud1<br>apo | Krimper Tud1-Ago3 | Krimper Tud2-AubR15me2 |
| --- | --- | --- | --- |
| **Data collection** |  |  |  |
| Beamline | SSRF-BL19U1 | SSRF-BL19U1 | SSRF-BL19U1 |
| Space group | $P2_12_12_1$ | $P3_1$ | $P2_12_12_1$ |
| Wavelength (Å) | 0.9792 | 0.9792 | 0.9792 |
| Cell dimensions (Å) |  |  |  |
| *a* | 60.5 | 88.5 | 97.1 |
| *b* | 87.7 | 88.5 | 101.4 |
| *c* | 90.2 | 181.7 | 191.3 |
| Resolution (Å) | 50.0-2.1 | 50.0-2.4 | 50.0-2.7 |
|  | (2.18-2.10)<sup>a</sup> | (2.49-2.40) | (2.80-2.70) |
| $R_{\text{merge}}$ | 0.134 (1.029) | 0.094 (0.891) | 0.114 (0.511) |
| $I / \sigma I$ | 12.5 (1.4) | 9.0 (1.25) | 12.5 (1.53) |
| Completeness (%) | 99.8 (100.0) | 99.3 (98.6) | 95.5 (75.1) |
| Redundancy | 5.4 (5.5) | 4.3 (4.3) | 5.1 (3.7) |
| CC1/2 | 0.727 | 0.872 | 0.897 |
| **Refinement** |  |  |  |
| $R_{\text{work}} / R_{\text{free}}$ | 0.204 / 0.222 | 0.234 / 0.259 | 0.243 / 0.291 |
| No. reflections | 28,403 | 61,721 | 49,822 |
| No. atoms | 3,274 | 7,519 | 12,195 |
| Protein | 3,062 | 6,899 | 11,794 |
| Peptide | - | 325 | 327 |
| Solvent | 212 | 295 | 74 |
| $B$-factors (Å<sup>2</sup>) | 39.3 | 56.4 | 58.1 |
| Protein | 39.1 | 56.0 | 57.9 |
| Peptide | - | 67.6 | 69.2 |
| Solvent | 42.8 | 53.3 | 50.7 |
| R.m.s. deviations |  |  |  |
| Bond lengths (Å) | 0.023 | 0.004 | 0.009 |
| Bond angles (°) | 1.771 | 0.743 | 1.237 |

<sup>a</sup> Highest-resolution shell is shown in parentheses.

