## Supplemental Table 2 for "Binding of guide piRNA triggers methylation of the unstructured N-terminal region of Aub leading to assembly of the piRNA amplification complex"

**Table S2. Oligo sequences.**

| **qPCR primers** | |
| --- | --- |
| HeT-A-F | CGCGCGGAACCCATCTTCAGA |
| HeT-A-R | CGCCGCAGTCGTTTGGTGAGT |
| ZAM-F | ACTTGACCTGGATACACTCACAAC |
| ZAM-R | GAGTATTACGGCGACTAGGGATAC |
| Burdock-F | AGGGAAATATTTGGCCATCC |
| Burdock-R | TTTTGGCCCTGTAAACCTTG |
| TAHRE-F | CTGTTGCACAAAGCCAAGAA |
| TAHRE-R | GTTGGTAATGTTCGCGTCCT |
| RP49-F | CCGCTTCAAGGGACAGTATCT |
| RP49-R | ATCTCGCCGCAGTAAACG |
| **RNA oligos** |  |
| 26nt ssRNA | rUrCrGrArArGrUrArUrUrCrCrGrCrGrUrArCrGrUrGrArUrGrUrU |
| 30nt ssRNA (size marker) | rCrCrArUrCrGrArUrArArArArGrUrUrUrArArArCrGrArGrCrUrUrCrCrCrG |
| 42nt ssRNA (size marker) | rCrCrArUrCrCrArUrCrGrArUrArArArArGrUrUrUrArArArCrGrArGrCrUrUrCrCrCrGrCrGrUrArCrGrGrA |
